# A Physics-Informed Neural Network Surrogate for Patient-Specific Hepatic Arterial Hemodynamics in Yttrium-90 Radioembolization: Network Architecture, Boundary-Condition Enforcement, and Data Efficiency

**DOI:** 10.64898/2026.09.26.754555

**Authors:** Erfan Taatizadeh, Nisanth Kumar Panneer Selvam, Emilie Roncali

## Abstract

Trans-arterial radioembolization using yttrium-90 (Y-90) microspheres treats unresectable liver cancer with radiation. The tumor-to-parenchyma dose ratio is mainly governed by the patient-specific hepatic arterial flow distribution, which transports the microspheres. Even though computational fluid dynamics (CFD) simulation can predict this flow distribution, its high computational cost and time limit intraprocedural application. To overcome this computational limitation, a physics-informed neural network (PINN) surrogate was developed to provide rapid predictions of flow distribution in a patient-specific hepatic artery model. The surrogate maps spatial coordinates to the velocity vector 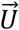 with Cartesian components *u, v, w* and the static pressure *p*. Reference velocity and pressure fields came from a converged finite-element CFD solve of the same geometry at about 1.05 × 10^5^ nodes; a controlled fraction was used to supervise the training while the remainder was held out for validation. Using the NVIDIA PhysicsNeMo framework, we compared four coordinate networks (MLP, Modified Fourier, Multi-Scale Fourier and SIREN) under soft and hard inlet enforcement across a supervised data sweep (data fraction *df* = 0 to 0.20) and repeated *df* = 0.20 at a matched 300,000-step budget. Predicted flow splits were propagated through an established Y-90 dose-kernel pipeline. Accuracy is governed primarily by the supervised-data fraction, not the architecture: at *df* ≥ 0.10 the backbones agree to within one to two *R*^2^ points, and *R*^2^(*u*) rises from ≈ 0 under pure physics (*df* = 0) to ≈ 0.95 by *df* = 0.01 for the best backbones, saturating near 0.97–0.99 by *df* = 0.20. At a matched 300,000-step budget the Multi-Scale Fourier network leads (*R*^2^(*u*) = 0.978) ahead of SIREN (*R*^2^(*u*) = 0.976) and the MLP (*R*^2^(*u*) = 0.968); the best value seen (*R*^2^(*u*) = 0.985, Modified Fourier) needed 1.8 times that budget. On the *df* = 0.20 checkpoint the per-outlet flow split matches CFD (*r* = 0.999, *L*^1^ = 4.4 percentage points) and the derived dosimetry agrees to within 2.4% for perfused volume, mean dose and every isodose volume. At *df* = 0 training collapses toward the trivial near-zero state, which satisfies continuity, momentum and the zero-pressure outlets exactly and is ruled out only by the inlet and data terms; this, not architecture choice, is the limiting failure mode. Once trained, the network evaluates the full 105,284-node field in about 2 s on a four-core CPU. The supervised-data fraction, with a practical floor between *df* = 0.05 and 0.10, is the primary design lever for such hepatic-arterial PINN surrogates.

## 1. Introduction

Liver cancer is among the leading causes of cancer death worldwide. For unresectable hepatocellular carcinoma and metastatic liver disease, trans-arterial radioembolization (TARE) with Y-90 microspheres delivers localized β-radiation through the hepatic arterial supply that preferentially perfuses tumors. The therapeutic ratio, dose to tumor versus dose to healthy liver, is determined by where microspheres lodge in the arterial tree, which in turn depends on patient-specific arterial tree geometry, catheter position, inlet flow waveform, and the downstream resistance that sets the flow split among branches [1], [2], [3], [4], [5]. The absorbed dose results from that flow field in two steps: microspheres are partitioned among the outlet branches in proportion to their flow share and tracked to where they lodge, then the resulting activity map is convolved with a Y-90 dose-point kernel to give the three-dimensional absorbed dose [6]. Pre-treatment planning currently relies on technetium-99m macroaggregated albumin (Tc-99m MAA) imaged with single-photon emission computed tomography (SPECT) and on vascular mapping with contrast-enhanced CT (CECT) and cone-beam CT (CBCT) by the interventional radiologist [1]. Tc-99m MAA SPECT has a spatial resolution of about 12 mm or coarser and does not consistently predict the Y-90 microsphere distribution [6], and neither imaging method can evaluate an alternative catheter position without a new injection. Patient-specific CFD, performed on hepatic arterial geometries segmented from CBCT or CECT, complements this workflow: it predicts microsphere transport and absorbed dose, and its predicted dose distribution was consistent with post-treatment Y-90 PET/CT [6]. Crucially, the predicted hepatic arterial flow distribution, and therefore the dose delivered, is highly sensitive to the outlet boundary conditions that represent the distal vasculature, which is the single largest modeling lever in hepatic TARE simulations [2]. A high-fidelity CFD solution of a patient hepatic arterial tree (order 10^6^–10^7^ elements, dozens of outlets) must be solved on commercial finite-element or finite-volume solvers for every new geometry and boundary condition [2], [6]; for the present tree a single solve takes hours (Section 2.6). This single-shot cost prevents two clinically valuable applications: real-time comparison of candidate catheter positions during a planning session, and intra-procedural re-planning when contrast injection reveals unanticipated flow behavior, such as reflux into extrahepatic branches, stasis in a branch, or a flow split that differs from the planning study [3], [4], [5]. A surrogate that returns velocity and pressure in milliseconds, while respecting the governing physics, would unlock both.

Physics-informed neural networks (PINNs) train a coordinate network to satisfy the governing partial differential equations (PDEs) and boundary conditions at sampled collocation points, with PDE residuals evaluated by automatic differentiation and optionally augmented with sparse data [7], [8]. Unlike purely data-driven surrogates, PINNs embed conservation laws directly and can operate with little or no labeled data; unlike operator-learning approaches (e.g., DeepONet [9]), a single-geometry PINN requires no library of pre-computed solutions and is the natural first step toward clinical deployment. PINNs have been applied across cardiovascular flows: reconstructing arterial pressure from 4D-flow MRI [10], improving cerebral hemodynamics from sparse Doppler and angiographic data [11], estimating three-element Windkessel outlet parameters from synthetic phase-contrast MRI [12], tracking the evolving false lumen of a dissected aorta in synthetic CFD data via transfer learning [13], and predicting 4D hemodynamics conditioned on vascular morphology [14]. Several seminal reviews of PINNs applied to fluid mechanics and biomedical engineering survey these advances and summarize open problems [7], [8], [15]. This paper focuses solely on PINNs for biofluid flow modeling. Figure 1 contrasts the conventional CFD and PINN pathways for this application.

**Figure 1.**
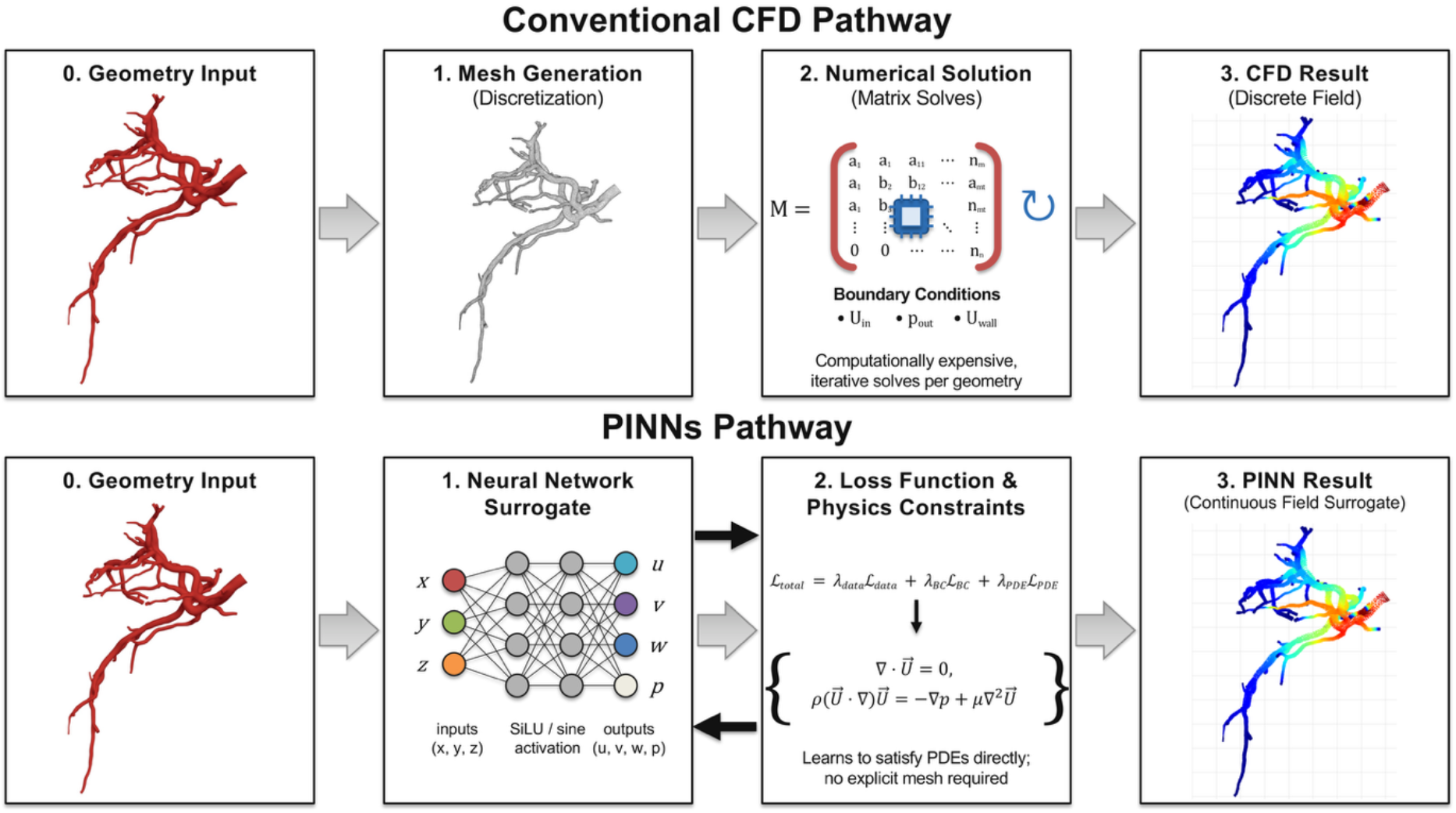
Conceptual comparison of the conventional CFD pathway and the physics-informed neural network (PINN) pathway for patient-specific hepatic arterial hemodynamics. The CFD pathway requires manual meshing and an expensive iterative computation for every new geometry; the PINN learns a continuous coordinate-to-field surrogate (*x, y, z*) ↦ (*u, v, w, p*), where (x, y, z) are spatial coordinates, (u, v, w) the Cartesian velocity components and p the static pressure, by minimizing a combined data and physics loss and, once trained, evaluates the whole field in seconds for the geometry and boundary conditions it was trained on.

Three difficulties dominate practical PINN training on complex geometries. First, soft-constrained PINNs are prone to optimization failure modes in stiff or convection-dominated regimes [16]; for steady incompressible flow, the constant state 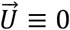, where 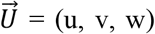 is the velocity field, with uniform pressure satisfies continuity, momentum, and no-slip exactly, and is a strong “trivial-solution” attractor when the data and inlet terms are weak [17]. The standard remedy is to supplement the physics residuals with a limited set of CFD samples, giving a hybrid, data-assisted PINN rather than a purely data-free one. Second, the volume of supervised data required for reliable convergence on multi-outlet vasculature is poorly characterized. Third, the choice of coordinate-network architecture strongly affects accuracy: standard multilayer perceptrons (MLPs) exhibit spectral bias toward low frequencies [18], which sinusoidal-representation networks (SIREN) [19], random Fourier-feature embeddings [18], and multiplicative-filter networks [20] are designed to overcome, and prior work on three-dimensional blood flow found architecture choice to change accuracy [21]. These three issues interact: the architecture and the boundary-condition enforcement together determine how much supervised data is needed to avoid the trivial attractor.

We make four contributions. (i) A working patient-specific PINN at clinically relevant Reynolds number for the hepatic arterial tree extracted from CBCT with 46 outlets, reaching *R*^2^ ≥ 0.96 for every velocity component, with full-field inference in about 2 s on a CPU. (ii) A comparison of four coordinate architectures matched on depth, width, data fraction and training budget. (iii) A supervised-data sweep showing that accuracy is governed primarily by the data fraction, with a practical floor between *df* = 0.05 and 0.10 and a sharp cliff at *df* = 0, plus an analytical and empirical characterization of the trivial-solution attractor that dominates there. (iv) An end-to-end Y-90 dosimetry evaluation showing that the surrogate preserves clinical dose endpoints to within a few percent.

## 2. Methods

The complete pipeline has three stages: a patient-specific geometry is solved once with CFD to provide reference fields, a coordinate network is trained against a fraction of those fields and the governing equations, and the resulting flow split is propagated through a Y-90 transport and dosimetry chain to an absorbed-dose map. Sections 2.1 to 2.5 set up the PINN pathway of Figure 1: the patient geometry, the governing equations, the network architectures, the boundary conditions, and the loss and training. Section 2.6 describes the COMSOL reference data and the accuracy metrics, and Section 2.7 the dose chain of Figure 2.

**Figure 2.**
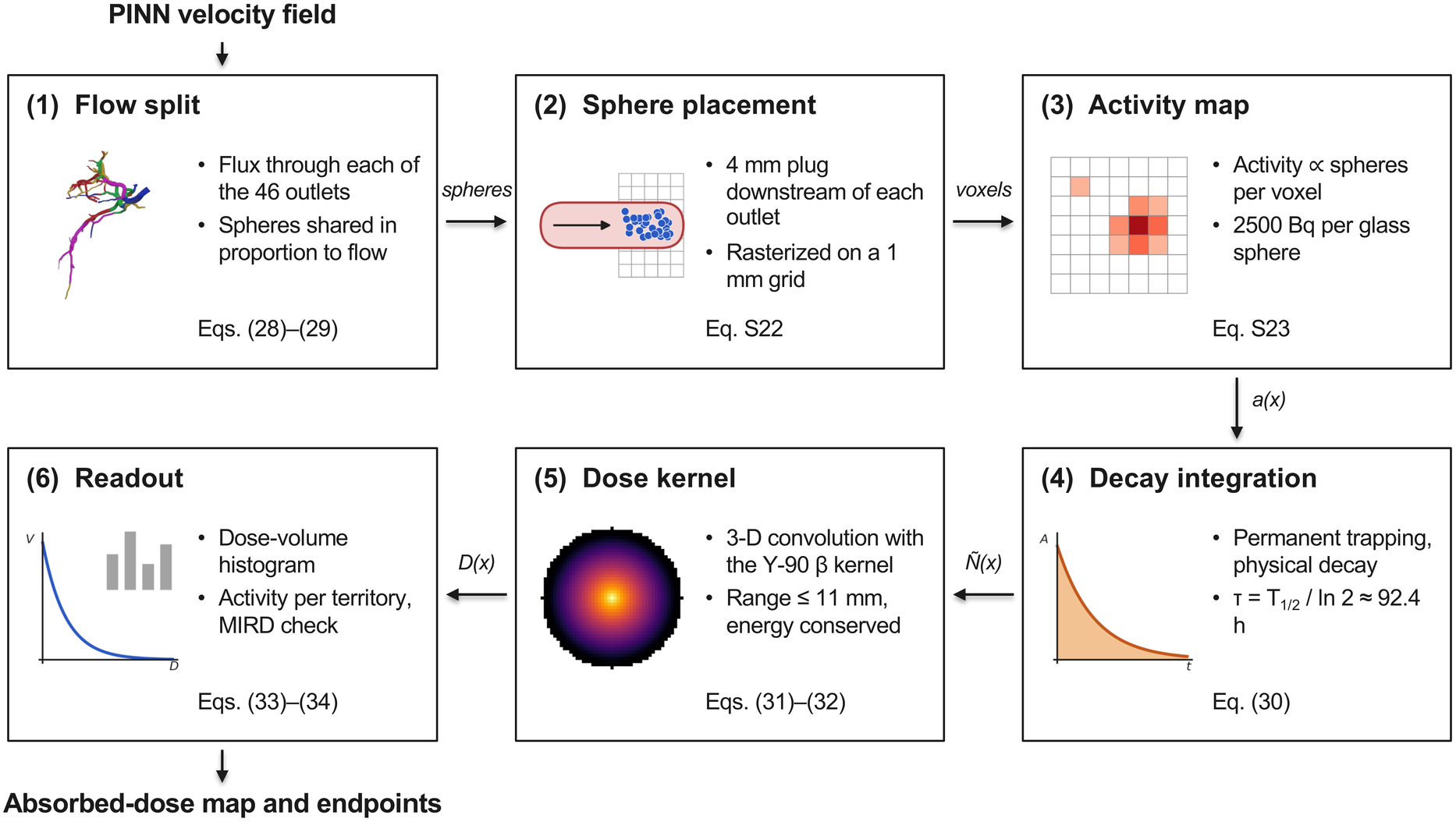
Flow-to-dose chain that turns the surrogate velocity field into a three-dimensional absorbed-dose map. Stage 1 converts the outlet fluxes into non-negative flow fractions and partitions the injected microspheres; stage 2 places each outlet’s spheres in a 4 mm cylindrical plug downstream of the outlet, rasterized on a 1 mm voxel grid, tagged by liver segment; stages 3 and 4 convert sphere counts into an activity map and then a time-integrated decay map; stage 5 convolves that map with the energy-conserving Y-90 β dose-point kernel; stage 6 reads out the dose-volume histogram, the per-segment activity shares and the whole-organ MIRD cross-check. Arrow labels give the quantity passed between stages; the tag in each box gives the governing equation in the main text or in Supplementary Note S1.

Figure 2 lays out the complete chain as an ordered pipeline, from the surrogate velocity field to the dose endpoints, with the governing relation for each stage marked. Apart from the random placement of spheres within each outlet cylinder, the chain is deterministic; besides the flow split, its only inputs are the injected activity, the specific activity of the spheres, the 4 mm lodging length and the dose-point kernel, none of which was tuned.

### 2.1 Patient geometry and computational domain

The hepatic arterial geometry was segmented from a contrast-enhanced cone-beam CT and coneq]verted to a watertight stereolithography (STL) surface after smoothing. The final mesh comprises 49,504 surface triangles with a single inlet and 46 distal outlets; the lumen volume is approximately 2.4 × 10^3^ mm^3^ and the inlet diameter is 4.57 mm (Table 1). For training, the geometry is rigidly rotated once so that the inlet lies in the *z* = 0 plane with inward normal +*z* and centroid at the origin; this alignment makes the inlet velocity profile a function of the in-plane radius alone and is a prerequisite for the hard inlet boundary condition. Rigid rotation leaves the physics invariant. Synthetic hepatic trees constructed via constrained constructive optimization [22] offer a route to scale this pipeline to many geometries in future work. The segmented lumen surface that defines the PINN domain is shown in Figure 3. The same tree is shown as a branching diagram in Figure 4, which makes the interdigitation of the territories explicit: geodesic tracing from the inlet to each outlet cap yields 43 bifurcations at depth levels of 2 to 14, and no ordering of the subtrees collects every territory into a single contiguous run.

**Table 1.**
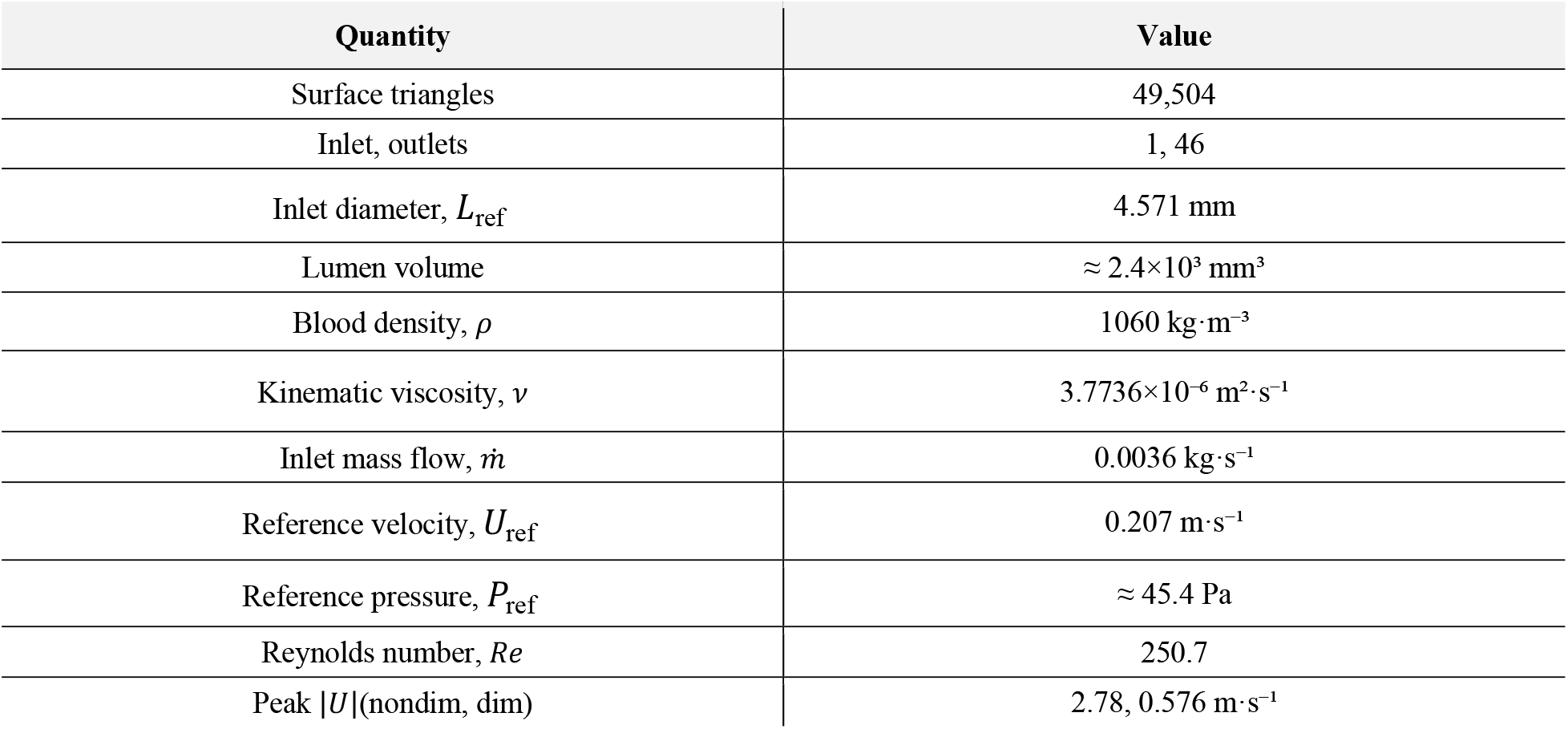
Geometry and physical parameters of the hepatic arterial model.

| Quantity | Value |
| --- | --- |
| Surface triangles | 49,504 |
| Inlet, outlets | 1, 46 |
| Inlet diameter, $L_{\text{ref}}$ | 4.571 mm |
| Lumen volume | $\approx 2.4 \times 10^3 \text{ mm}^3$ |
| Blood density, $\rho$ | $1060 \text{ kg} \cdot \text{m}^{-3}$ |
| Kinematic viscosity, $\nu$ | $3.7736 \times 10^{-6} \text{ m}^2 \cdot \text{s}^{-1}$ |
| Inlet mass flow, $\dot{m}$ | $0.0036 \text{ kg} \cdot \text{s}^{-1}$ |
| Reference velocity, $U_{\text{ref}}$ | $0.207 \text{ m} \cdot \text{s}^{-1}$ |
| Reference pressure, $P_{\text{ref}}$ | $\approx 45.4 \text{ Pa}$ |
| Reynolds number, $Re$ | 250.7 |
| Peak $\vert U\vert $(nondim, dim) | 2.78, $0.576 \text{ m} \cdot \text{s}^{-1}$ |

**Figure 3.**
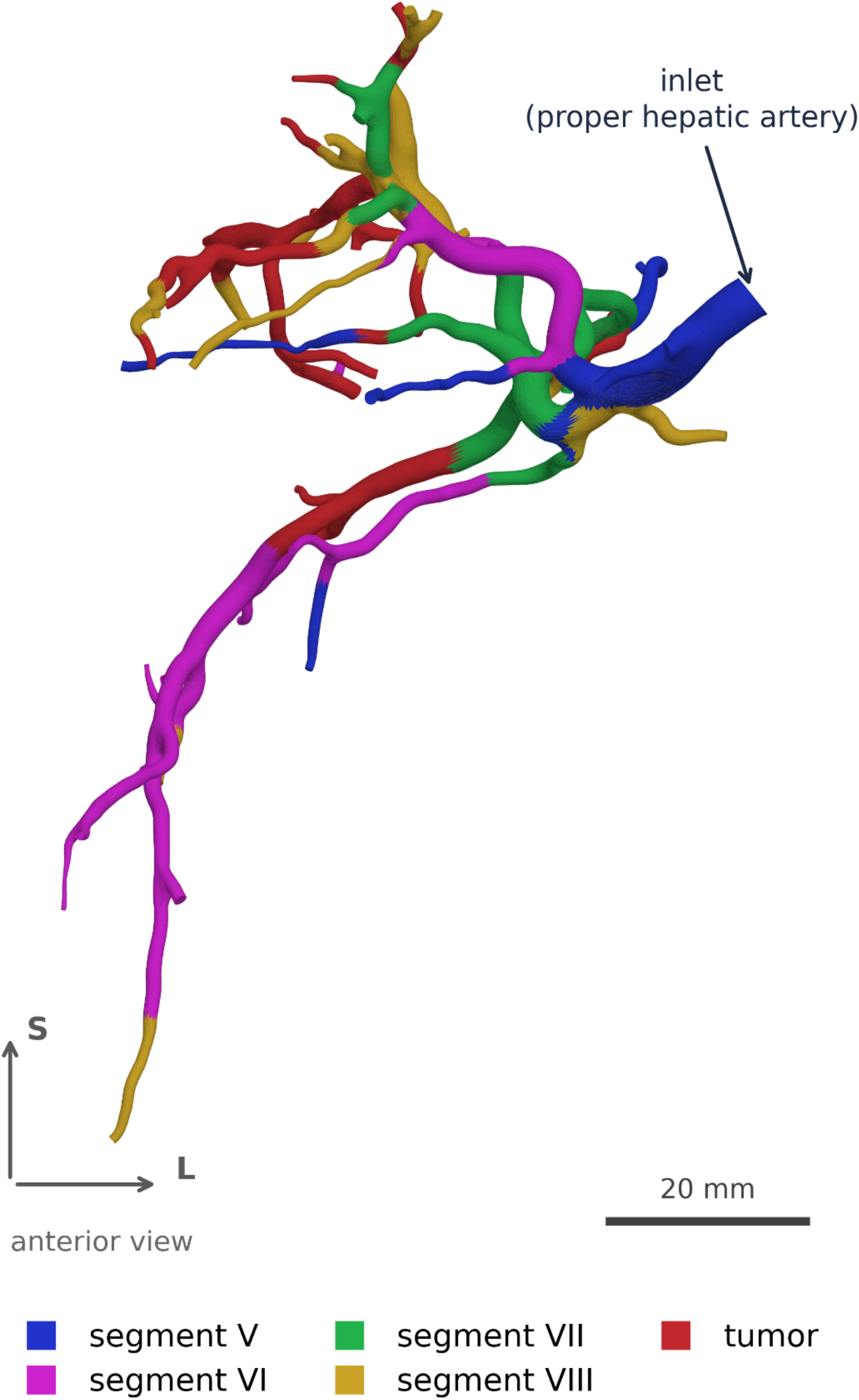
Patient-specific hepatic arterial geometry segmented from contrast-enhanced cone-beam CT (CBCT; one inlet, 46 distal outlets; 49,504 surface triangles), shown as the watertight lumen surface that defines the PINN domain Ω and colored by perfusion territory. Each facet, including those of the shared proximal trunk, takes the territory of the nearest outlet measured along the lumen surface. Anterior view in the CT frame: superior upward, the patient’s right on the viewer’s left, so segment VI descends to the lower left and the tumor territory lies upper left. Scale bar 20 mm. The solver instead works on a rigidly rotated copy with the inlet at *z* = 0.

**Figure 4.**
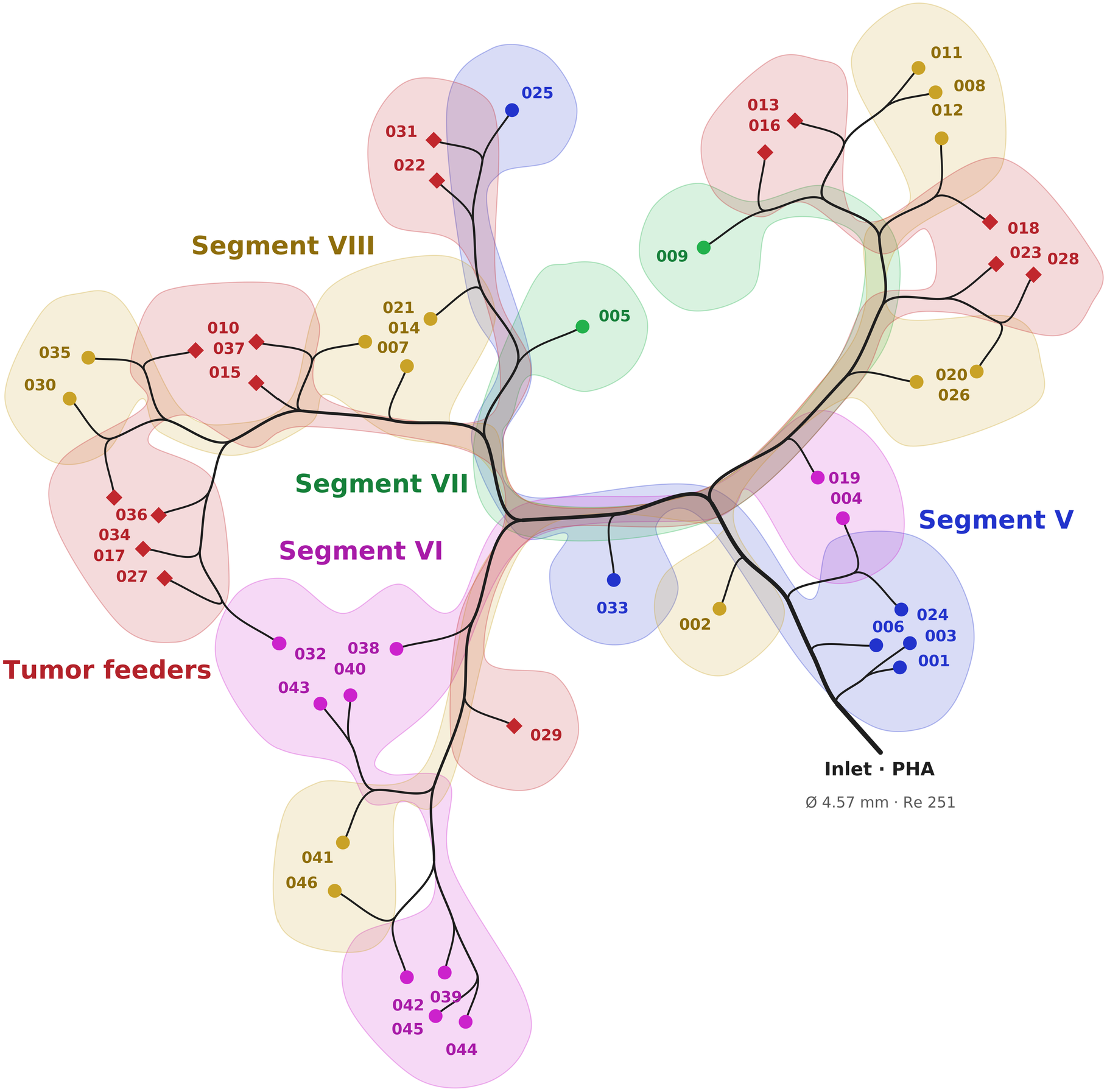
Branching topology of the same geometry, drawn as an unrooted dendrogram in the anterior view and colors of Figure 3, so that each territory lies on the same side as in the anatomy: the inlet enters at lower right and each tip is one of the 46 outlet caps. Branch length is proportional to centerline distance between bifurcations (compressed for legibility) and thickness to the number of outlets downstream. Each perfusion territory is shaded as one enclosed area; where a territory reaches the tree in more than one place the shading narrows and follows the branches that join those parts. Even with the subtree order at every bifurcation chosen by exact optimization over all flips, the 46 outlets break into 19 same-territory runs, the fewest this anatomy allows. Segment outlets are drawn as discs and tumor-feeding outlets as diamonds, so the two read apart without relying on color alone.

### 2.2 Governing equations and nondimensionalization

Blood is modeled as an incompressible, Newtonian fluid in the steady regime. The flow domain is the lumen Ω ⊂ ℝ^3^, with boundary ∂Ω = Γ_*in*_ ∪ Γ_*out*_ ∪ Γ_*wall*_, where Γ_*in*_ is the inlet cap, Γ_*out*_ the 46 outlet caps and Γ_*wall*_ the vessel wall. The dimensional steady Navier-Stokes momentum and continuity equations are

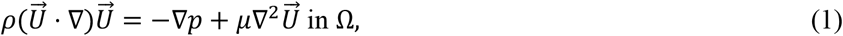

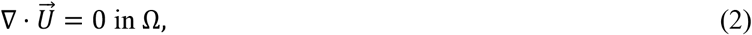

where 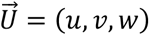 is the velocity field, p the static pressure, ρ the blood density, and µ = ρ*v* the dynamic viscosity. Equations (1)–(2) are closed by the boundary conditions: a prescribed inlet velocity profile on Γ_*in*_ zero pressure on the 46 distal outlets Γ_*out*_, and the no-slip condition 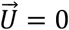 on the vessel wall Γ_*wall*_. The steady formulation represents the time-averaged operating point used in current dosimetry workflows; the unsteady (pulsatile) extension is left to future work.

Fields and coordinates are nondimensionalized by the inlet diameter *L*_*ref*_ and the bulk inlet velocity 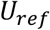, with the pressure scaled by the dynamic pressure 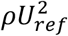:

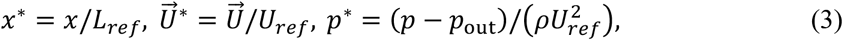

where *p*_*out*_ is the outlet reference pressure (zero here), the reference velocity follows from the prescribed inlet mass flow 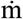 and the inlet cross-sectional area *A*_in_, and the Reynolds number is defined in the usual way:

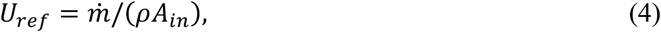

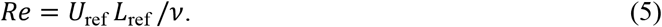

Substituting (3)–(5) into (1)–(2) yields the nondimensional system solved by the network. An asterisk marks a nondimensional quantity throughout, in the equations and in every figure axis; gradients are taken with respect to the nondimensional coordinates:

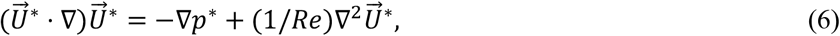

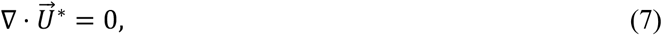

so that the single similarity parameter is Re. With the values in Table 1, *Re* ≈ 251 and the simulation (COMSOL) peak velocity magnitude is |*U*|_*max*_ ≈ 2.78 (nondimensional; 0.576 m · s^−1^), located near the proximal bifurcation. The PINN was trained with ρ = 1060 kg · m^−3^.

### 2.3 Coordinate-network architectures

Every architecture realizes a coordinate-to-field map with trainable parameters θ,

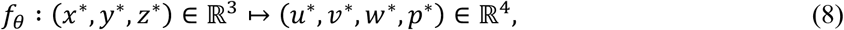

and shares a depth of six hidden layers of width 256; the four backbones (Table 2) differ mainly in their input encoding and activation. The baseline multilayer perceptron (MLP) is a fully connected network with the smooth SiLU activation, and every linear layer is reparameterized by weight normalization, which decouples each weight vector’s magnitude from its direction [23].

**Table 2.**
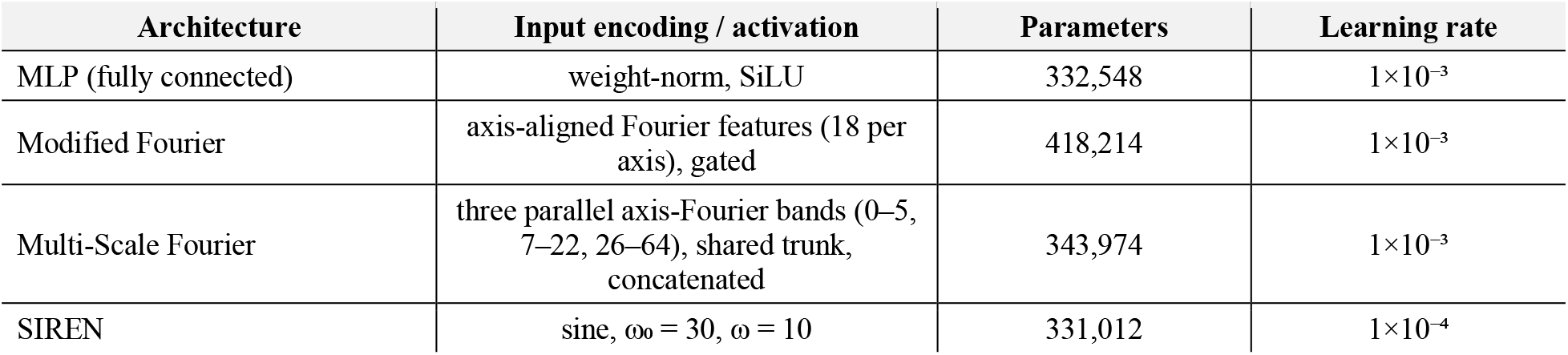
Coordinate-network architectures compared (all six hidden layers × 256 units). Parameter counts measured from the trained checkpoints.

| Architecture | Input encoding / activation | Parameters | Learning rate |
| --- | --- | --- | --- |
| MLP (fully connected) | weight-norm, SiLU | 332,548 | $1 \times 10^{-3}$ |
| Modified Fourier | axis-aligned Fourier features (18 per axis), gated | 418,214 | $1 \times 10^{-3}$ |
| Multi-Scale Fourier | three parallel axis-Fourier bands (0–5, 7–22, 26–64), shared trunk, concatenated | 343,974 | $1 \times 10^{-3}$ |
| SIREN | sine, $\omega_0 = 30$, $\omega = 10$ | 331,012 | $1 \times 10^{-4}$ |

Standard MLPs exhibit a spectral bias toward low-frequency functions [18], which the remaining three architectures counter in different ways. The Modified Fourier network first lifts the input through an axis-aligned Fourier-feature embedding with 18 frequencies per axis, then mixes two parallel encoders through per-layer multiplicative gates that reinject the encoding at every depth.

The Multi-Scale Fourier network instead applies *M* = 3 independent axis-aligned Fourier encodings, one per frequency band, to the same input coordinates in parallel, rather than forming the output as a product of filtered inputs in the manner of multiplicative-filter networks [20]. The three bands use the axis frequency sets *F*_1_ = {0, … , 5}, *F*_2_ = {7, … , 22} and *F*_3_ = {26, … , 64}; a single six-layer trunk whose weights are shared across all three bands processes each encoding, and the band outputs are concatenated and mapped to the field variables by one final linear layer, so the encodings are combined additively at the output rather than multiplicatively at every depth. Finally, the sinusoidal-representation network (SIREN) composes sine-activated layers with the frequency-scaled initialization of Sitzmann et al. [19].

Because any derivative of a SIREN is itself a SIREN, derivative supervision through the PDE residual remains well conditioned. The four backbones are specified formally in Supplementary Note S1 (Eqs. S1– S11). Parameter counts (Table 2) were verified directly from the trained checkpoints; Figure 5 summarizes the network and its composite training objective, and Figure 6 compares the four backbone topologies side by side.

**Figure 5.**
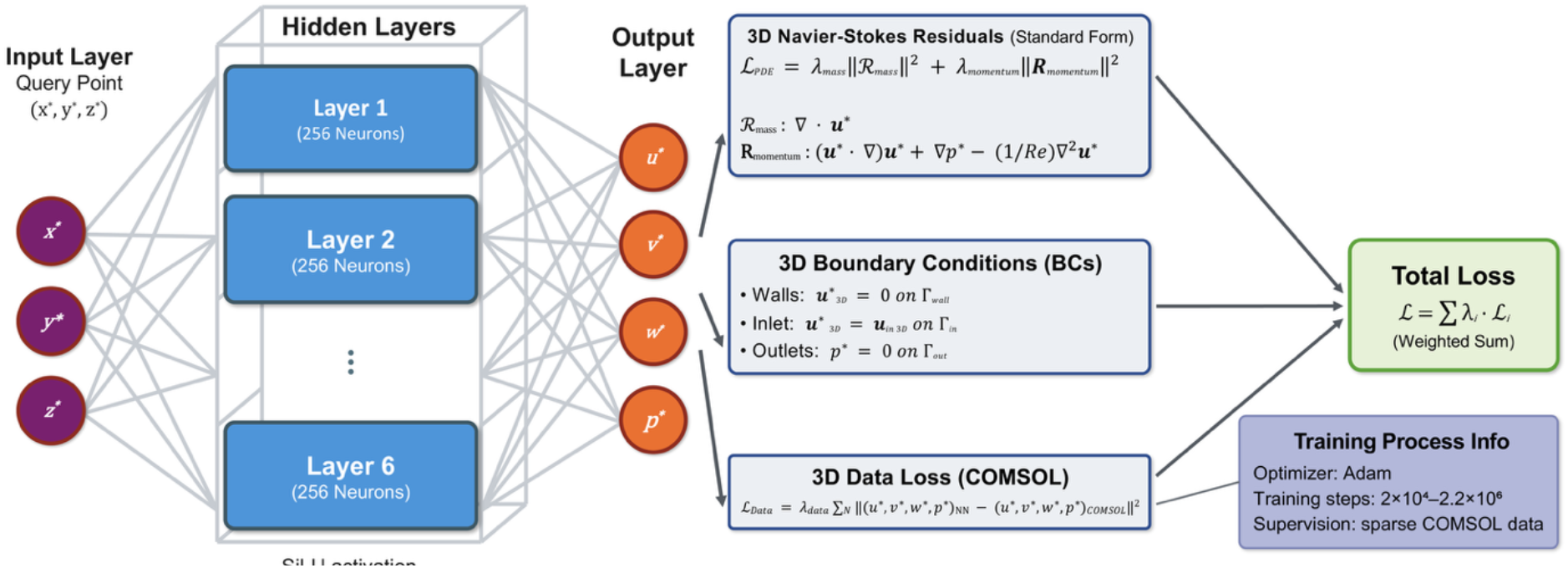
Architecture and loss formulation of the PINN. A coordinate query point (x, y, z) is mapped through six hidden layers of width 256 (SiLU activation; sine for SIREN) to the velocity components and pressure (u, v, w, p). Training minimizes a weighted sum of the 3D Navier-Stokes residuals (mass and momentum), the multi-surface boundary conditions (no-slip walls, prescribed inlet profile, zero-pressure outlets), and a sparse COMSOL data term, optimized with Adam (2×10^4^ to 2.2×10^6^ steps; 3×10^5^ in the step-matched comparison).

**Figure 6.**
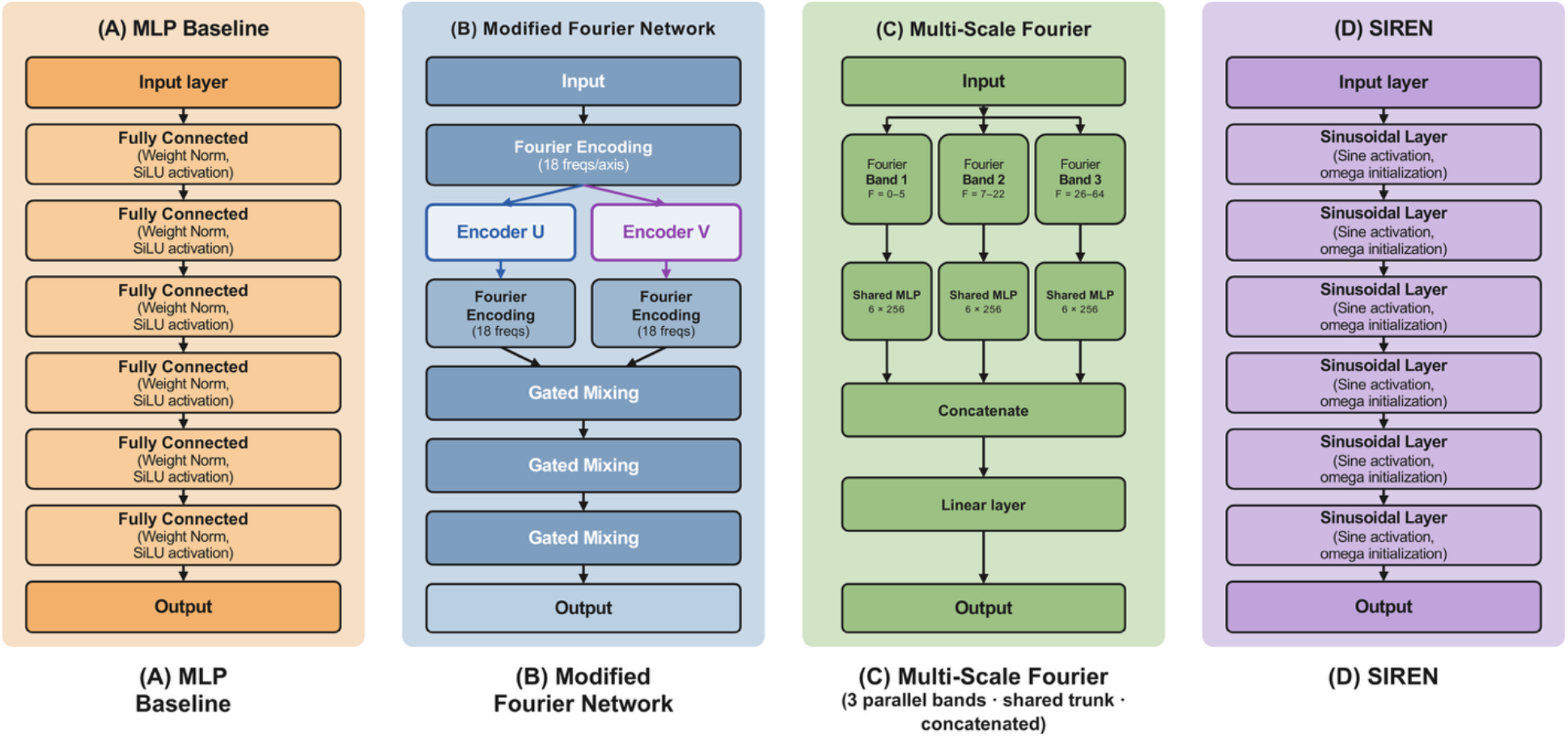
The four backbones compared at matched depth (six hidden layers) and width (256 units): (A) a weight-normalized MLP with SiLU activations; (B) a Modified Fourier network with a Fourier-feature embedding, two parallel encoders and per-layer gated mixing; (C) a Multi-Scale Fourier network applying three parallel axis-aligned Fourier encodings (bands 0– 5, 7–22, 26–64) to the same query point, propagating each through a shared trunk and concatenating at one final linear layer; and (D) a SIREN with sinusoidal activations and variance-preserving initialization.

### 2.4 Boundary conditions and enforcement

The inlet flow is prescribed as an axial velocity profile aligned with the inward normal, with a power-law profile whose exponent *n* is auto-fitted to the near-inlet COMSOL data (here *n* = 2.14, close to parabolic: its peak-to-mean velocity ratio is 1.94, against 2 for Poiseuille flow; the data-free runs, which have no reference data to fit, used n = 1.2), normalized so that its area-average equals the bulk velocity *U*_*m*_. The influence of the delivery catheter on such inlet boundary conditions in vascular CFD is modeled by Taebi et al. [24].

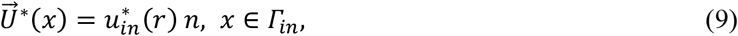

where the power-law profile is given in Supplementary Note S1 (Eqs. S12–S13) and r is the in-plane radius. Global mass conservation and the absence of swirl are imposed through an integral flux constraint and two tangential conditions on the inlet plane (Eqs. S14–S15). The 46 distal outlets impose a uniform zero pressure, and the wall is no-slip:

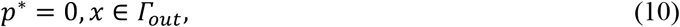

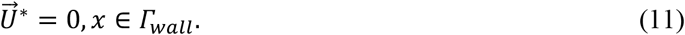

Equations (9)–(11), together with Eqs. S12–S15, define the soft formulation, in which each condition becomes a weighted penalty in the loss. In the hard formulation the inlet condition is instead built into the architecture. The geometry is rotated so that the inlet lies at *z* = 0 and the raw network outputs are blended toward the prescribed profile by a smooth hyperbolic-tangent gate of width δ (Supplementary Note S1, Eqs. S16–S18).

At the inlet plane the gate vanishes, so the output reproduces the prescribed profile for any network weights, while deep in the bulk the gate saturates to unity and the network is unconstrained. The inlet residual is therefore exactly zero by construction, and the inlet penalties are dropped. This ansatz follows the hard-constraint philosophy of data-free Navier-Stokes PINNs [25] and removes the failure mode in which the network satisfies the interior physics and the outlet condition while ignoring the inlet.

### 2.5 Loss formulation and training

The interior physics is enforced through the continuity and momentum residuals, evaluated pointwise by automatic differentiation of the network outputs,

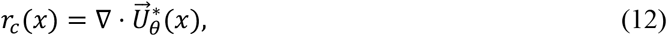

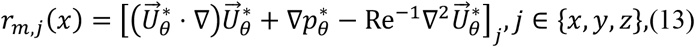

and the network is trained by minimizing a weighted composite objective function over the parameters θ,

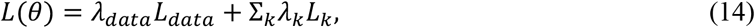

n which *k* runs over the residual and boundary terms, *k* ∈ {cont, mom, in, mdot, tan, out, wall}, and λ_*data*_ and λ_*k*_ are fixed weights. Each term is a mean-squared penalty evaluated on its own point set, defined term by term in Eqs. (15) to (22) below; the list after Eq. (22) states what each term enforces.

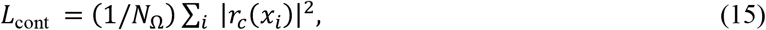

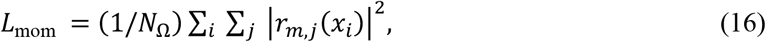

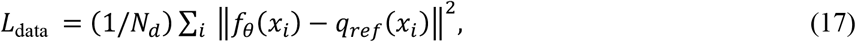

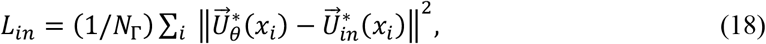

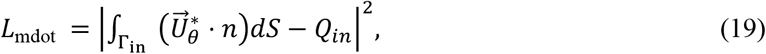

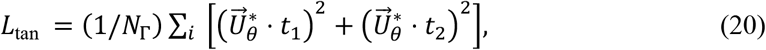

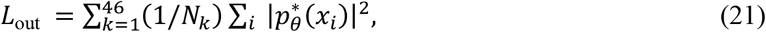

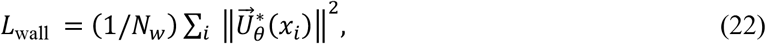

- *L*_cont_ (15): mass conservation, through the continuity residual at the interior collocation points;
- *L*_mom_ (16): momentum balance, through the three momentum residuals at the same points;
- *L*_data_ (17): agreement with the COMSOL velocity and pressure at the supervised points;
- *L*_*in*_ (18): the prescribed inlet velocity profile;
- *L*_mdot_ (19): the prescribed inlet flow rate, through the flux across the inlet;
- *L*_tan_ (20): zero tangential velocity at the inlet;
- *L*_out_ (21): zero pressure on the 46 outlet caps, each outlet weighted equally;
- *L*_wall_ (22): no-slip at the vessel wall.

The data term compares the network output with the COMSOL reference field on the supervised fraction only. Under hard inlet enforcement the inlet terms, Eqs. (18) to (20), are removed. We use the simple summation aggregator with fixed weights (λ_*data*_ = 5, λ_wall_ = 5, and unit weights on continuity, momentum, outlet pressure and the three inlet terms), deliberately avoiding adaptive re-weighting schemes [26] so that the observed data-efficiency behavior is attributable to the data fraction rather than to loss balancing. The minimizer is sought with the Adam optimizer under an exponentially decaying learning-rate schedule (Supplementary Note S1, Eqs. S19–S20). A lightweight per-constraint logging modification of the solver records each (constraint, variable) loss separately, exposing the trivial-solution signature that aggregated logging hides.

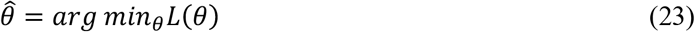

Training used NVIDIA PhysicsNeMo Sym (v2) [27] on a single NVIDIA RTX 5070 Ti GPU. Interior (800,000) and wall (80,000) collocation pools were pre-sampled once; per-step interior and wall batches were 10,000 points for the MLP and Modified Fourier networks, 5,100 for the Multi-Scale Fourier network (limited by GPU memory) and 30,000 for most SIREN runs, with 1,500 data and inlet points and 250 per outlet. Optimization used Adam with an exponential learning-rate decay (rate 0.95 every 20,000 steps); the initial rate was 10^−3^ for the Fourier-family and MLP networks and 10^−4^ for SIREN to control the ω-dependent gradient amplification of sinusoidal layers. Runs spanned 2×10^4^ to 2.2×10^6^ steps (hours to days of wall time). Stability of deep coordinate networks benefits from residual (skip) connections [28]; here weight normalization sufficed, as the networks are six layers deep at most and showed none of the gradient issues that motivate residual connections.

### 2.6 Reference data and accuracy metrics

Accuracy is reported as 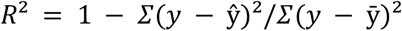 on held-out points, with the relative *L*^2^ error and, in physical units, RMSE, MAE and signed bias. The value the validator reports, 1 − (relative *L*^2^)^2^, which is used in the sweep tables, and the standard *R*^2^ agree to within 0.0012 for every field here, so the two are interchangeable; every value is quoted with its data fraction and step count. On the reference *df* = 0.20 model the RMSE is 0.0044, 0.0052 and 0.0060 m s^−1^ for *u, v* and *w* and 0.72 Pa for pressure against a peak speed of 0.576 m s^−1^, with signed biases below 5 × 10^−4^ m s^−1^ and 0.12 Pa.

Reference fields were computed in COMSOL Multiphysics (steady incompressible laminar flow) on a tetrahedral mesh, with the same power-law inlet, zero-pressure outlets and no-slip walls, and exported as volumetric (*u, v, w, p*) at ≈ 1.05 × 10^5^ points. A single solve takes hours, establishing the cost baseline the surrogate replaces.

To quantify supervised-data requirements, a fraction *df* of the COMSOL points (capped at 105,000, essentially the whole COMSOL export of 105,284 nodes, so that df is a fraction of the complete reference field) was used as a pointwise data term during training and the remainder held out for validation; *df* ∈

{0, 0.001, 0.005, 0.01, 0.05, 0.10, 0.20}. The sweep stops at *df* = 0.20 by design: the study asks how little supervision suffices, and stopping at 0.20 keeps 80% of the reference field, 84,000 points, for validation. Accuracy has also plateaued by then (Section 3.2), so larger fractions would add little. At *df* = 0 the network is data-free (physics and boundary terms only); a measurement-only validator was retained so that even non-converging runs can be scored against COMSOL.

Accuracy is reported on the held-out validation points for each field *q* ∈ {*u, v, w, p*} and for the velocity magnitude

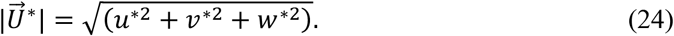

The primary metric is the coefficient of determination *R*^2^, complemented by the relative L^2^ error,

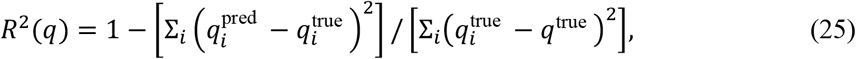

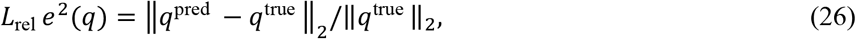

These are equivalent up to a factor *κ* that depends only on the ground truth, namely the ratio of the mean square of the true field to its variance (Supplementary Note S1, Eq. S21).

To check that the network does not smooth out the velocity peak, we additionally report the peak-velocity capture ratio

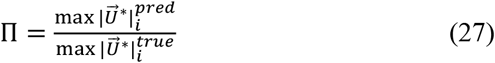

with Π = 1 indicating exact capture. All quantities were computed directly from the validator point clouds (*κ* ≈ 1.03,1.03,1.06 for *u, v, w* and ≈ 2.13 for *p*).

### 2.7 CFD-based Y-90 dosimetry

The validated velocity field drives a microsphere-transport and absorbed-dose calculation that follows the CFDose framework of Roncali and Taebi [1], [2], [6]. Before the transport calculation the surrogate velocity is returned to physical units by multiplying by the reference velocity, so every quantity from here on is dimensional. The volumetric flow through each outlet face Γ_*k*_ is the flux of that velocity through the face,

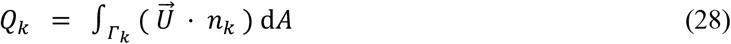

where *n*_*k*_ is the outward unit normal of the cap and *A*_*k*_ its area. Microspheres are partitioned among the outlets in proportion to their non-negative flow share,

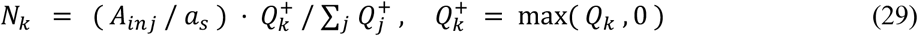

where *A*_*inj*_ is the injected activity and *a*_*s*_ the specific activity (2500 Bq for glass, 50 Bq for resin; in this work, we used glass spheres) [6]. Spheres lodge in a short cylinder of radius 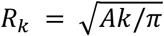 and 4 mm length downstream of each outlet, placed uniformly at random as point sources of activity (sphere size does not enter), and are accumulated on a 1 mm voxel grid as an activity map *a*(*x*); the cumulative number of β decays in a voxel is

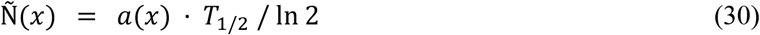

with the Y-90 physical half-life *T*_1/2_ = 64.05 h. The absorbed-dose map is obtained by convolving the decay map with a Y-90 dose-point kernel *K*(*r*),

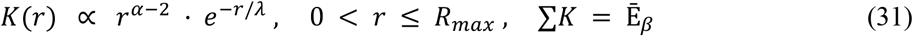

where α = 1.3 and λ = 1.1 mm are shape parameters of the Y-90 β kernel (Supplementary Figure S1), with maximum β range *R*_*max*_ = 11 mm and mean β energy per decay Ē_*N*_ = 0.9337 MeV, giving the per-voxel dose

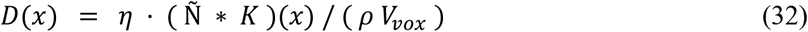

where ∗ denotes three-dimensional convolution, η the MeV-to-joule conversion, ρ the tissue density and *V*_*vox*_ the voxel volume. The per-segment activity share *f*_*s*_ and the cumulative dose-volume histogram *V*(*D*) follow directly,

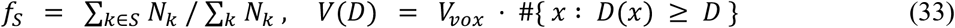

and the whole-organ dose is cross-checked against the standard MIRD estimate (*A* in GBq, *m* in kg) [6],

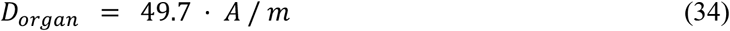

In Eq. (33), *V*(*D*) is the volume of tissue receiving at least a dose *D*: the perfused volume is the volume above 1 Gy, the mean perfused dose is the average dose over that volume, and *V*_50_ to *V*_400_ are the volumes receiving at least 50 to 400 Gy.

This pipeline follows the CFDose dosimetry of Roncali et al. [6] with the PINN surrogate substituted for the finite-element solver and the analytic kernel of Eq. (31) in place of their Monte Carlo kernel; the resulting endpoints are compared with COMSOL CFD simulation results in Section 3. The analytic kernel can be regenerated at any voxel size without a Monte Carlo run, and because the PINN and COMSOL dose maps share it, any kernel error cancels in their comparison.

The model treats microspheres as passive flow tracers, neglecting inertia, near-wall margination and catheter-jet effects; idealizes lodging as a 4 mm cylindrical plug at each outlet rather than at the arteriolar scale; assumes homogeneous water-equivalent tissue in the kernel convolution; and assumes permanent trapping, so that only physical decay governs the time integral. For a transient (pulsatile) PINN the outlet flux *Q*_*k*_(*t*) is cycle-averaged before forming the flow fractions; the decay integral τ is unaffected.

## 3. Results

### 3.1 Matched-budget comparison

At a matched supervised-data fraction and a matched training budget the four architectures are closely comparable, and accuracy is set primarily by the data fraction rather than the backbone. The most controlled test is the step-matched set at *df* = 0.20 with hard inlet enforcement and a matched ≈ 300,000-step budget (Table 3, Figure 7): the three step-matched backbones rank Multi-Scale Fourier, SIREN and MLP, with all four fields listed in Table 3, a spread of 0.010 in *R*^2^(*u*) at an equal step count. Switching the Multi-Scale Fourier run to soft enforcement increases *R*^2^(*u*) by 0.001, so the ordering is not an artifact of the inlet treatment. The Modified Fourier network attains the highest values (Table 4; Figures S6 and S7) but at 536,000 steps, and inside the matched budget it reaches 0.963; we report it as best observed rather than best at an equal step count.

**Table 3.** Step-matched architecture comparison at df = 0.20 and a common ≈ 300,000-step budget, the controlled test of the architecture effect. Validator R^2^ on held-out points at the final step; hard inlet enforcement unless noted. The Modified Fourier row is under-budget: its only df = 0.20 hard-BC run stopped at 100,000 steps, and its best result (0.985) came from a 536,000-step soft-BC run (Table 4).

| Architecture | BC | Steps | $R^2(u)$ | $R^2(v)$ | $R^2(w)$ | $R^2(p)$ | Note |
| --- | --- | --- | --- | --- | --- | --- | --- |
| Multi-Scale Fourier | hard | 300,000 | 0.978 | 0.960 | 0.981 | 1.000 | matched |
| SIREN | hard | 293,100 | 0.976 | 0.957 | 0.978 | 0.999 | matched |
| MLP | hard | 300,000 | 0.968 | 0.951 | 0.976 | 0.999 | matched |
| Multi-Scale Fourier | soft | 300,000 | 0.979 | 0.964 | 0.981 | 1.000 | matched, soft BC |
| Modified Fourier | hard | 100,000 | 0.963 | 0.941 | 0.974 | 0.999 | under-budget |

**Table 4.**
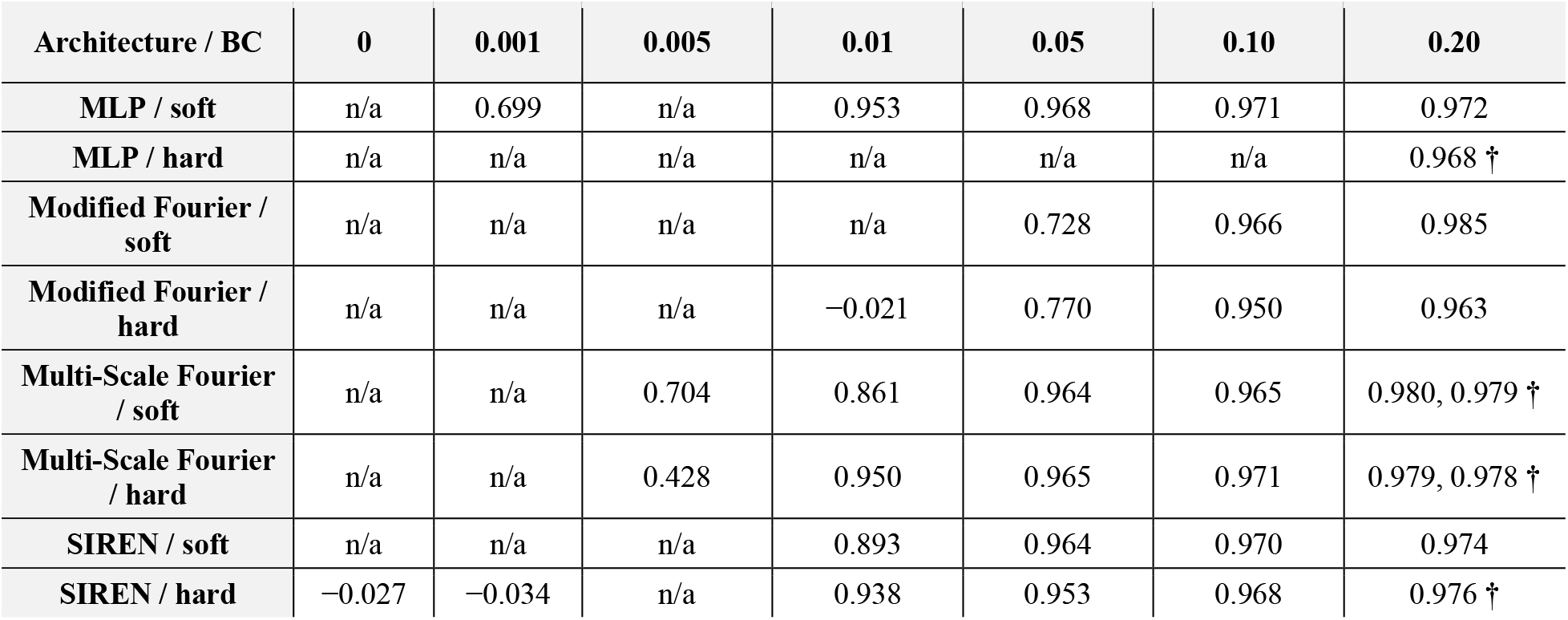
Validator *R*^2^(*u*) by architecture, inlet boundary condition and supervised-data fraction *df*, at the longest available budget for each cell. All runs except one data-free run share the same flow (*Re* ≈ 251); † marks the step-matched *df* = 0.20 runs of Table 3; where a cell has two budgets, both are given, the longest first. The *df* = 0 column shows the longest pure-physics SIREN run at this flow. The complete 41-run table is Supplementary Table S1.

**Figure 7.**
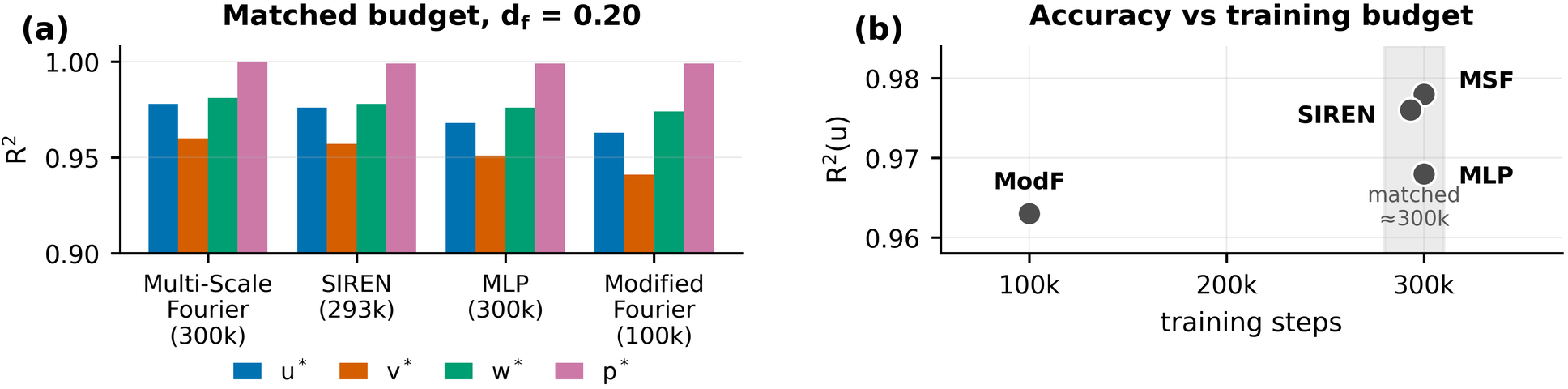
Step-matched architecture comparison at *df* = 0.20. (a) *R*^2^ for all four fields, by backbone, at a common ≈ 300,000-step budget; the Modified Fourier bars are from its only *df* = 0.20 hard-BC run, which stopped at 100,000 steps. (b) *R*^2^(*u*) against training budget, with the matched window shaded. At an equal step count, the three step-matched backbones span 0.010 in *R*^2^(*u*), that is about one *R*^2^ point.

In the rest of the results, the configuration with 20% of the full data set (*df* = 0.20) and the Multi-Scale Fourier model with soft inlet enforcement (300,000 steps) will serve as the reference configuration for comparing flow fields and dose distributions. Figure 8 shows the resulting velocity-magnitude and pressure fields beside the COMSOL reference, with mean absolute errors of 0.64% and 0.17% of the field range, and Figure 9 quantifies the same agreement on held-out points. Pressure is recovered almost exactly because the zero-pressure outlets anchor its scale, whereas the transverse velocity *v* is hardest to fit. The peak-velocity capture ratio of the reference model is Π = 1.001 (0.992 under hard enforcement), so the network does not smooth out the velocity peak. Once trained, it evaluates the full 105,284-node field in 2.2 s and the 46 outlet fluxes in 0.3 s on a four-core CPU, against hours for a CFD solve of the same geometry.

**Figure 8.**
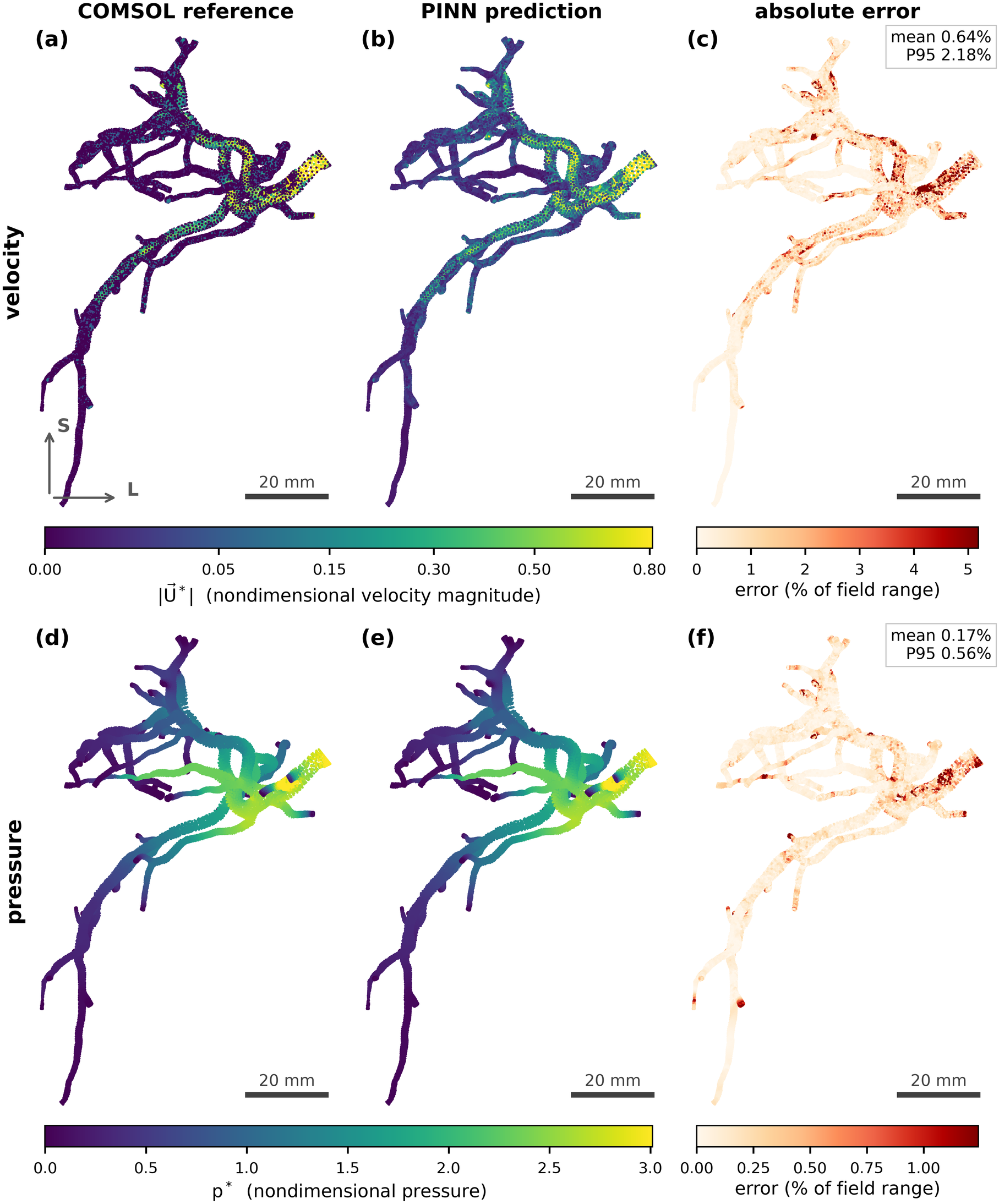
Velocity and pressure fields for the reference model (Multi-Scale Fourier, *df* = 0.20), shown in the anterior view of Figure 3. Top row, velocity magnitude: (a) COMSOL reference and (b) PINN prediction share one color scale; (c) absolute error as a percentage of the field range, concentrated around the proximal bifurcation, with scattered hot spots in small distal branches (mean 0.64%, P95 2.18%). Bottom row, pressure: (d) COMSOL reference, (e) PINN prediction, (f) absolute error. Pressure is recovered almost exactly (mean 0.17%, P95 0.56%) because the zero-pressure outlets anchor its scale; residual error sits near the inlet. Scale bar 20 mm.

**Figure 9.**
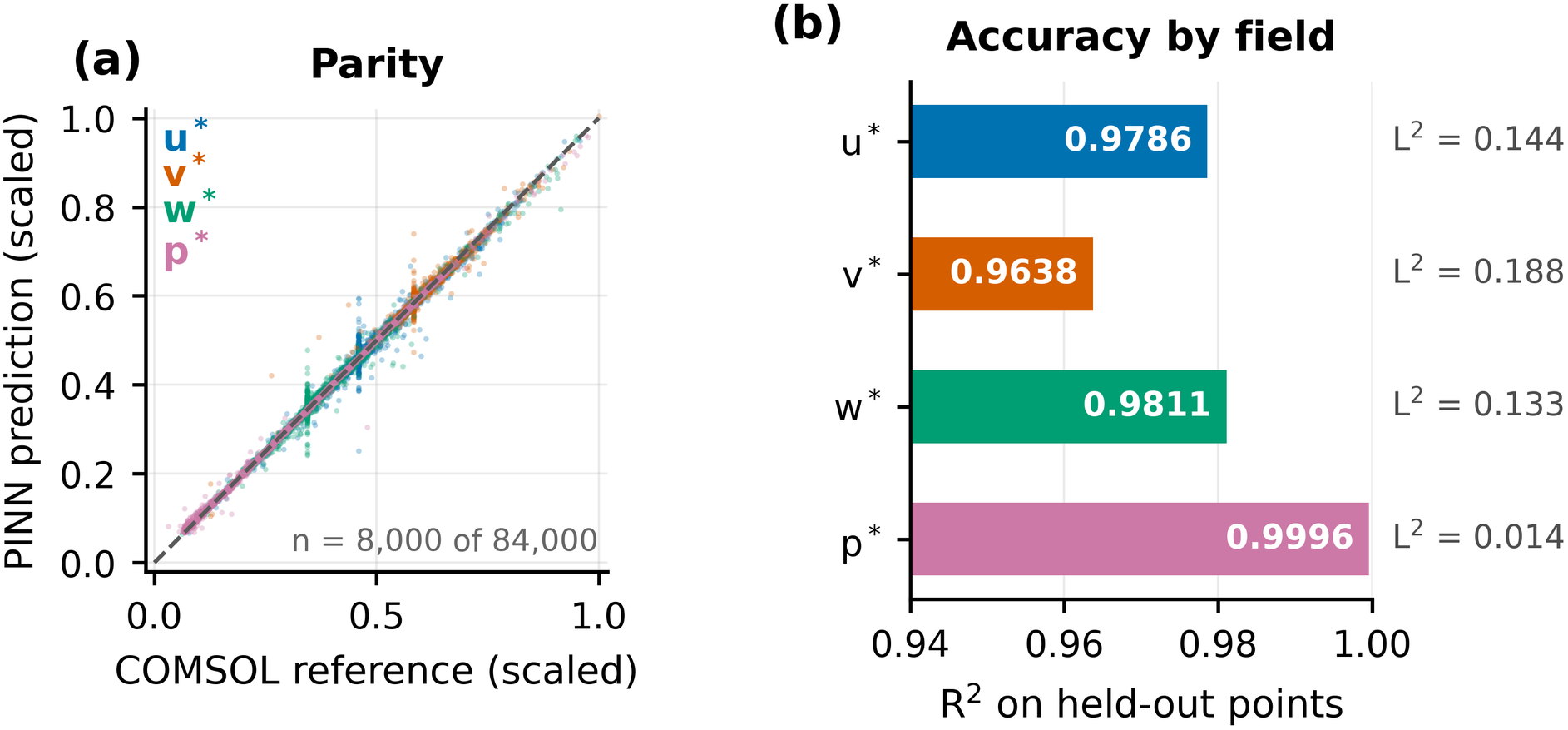
Accuracy of the reference model (Multi-Scale Fourier, *df* = 0.20, 300,000 steps) on held-out points. (a) Parity, with each field scaled to its own range so that no single variable dominates the axis. (b) *R*^2^ by field on a zoomed axis, with the relative *L*^2^ error printed alongside. Pressure is recovered almost exactly (*R*^2^ = 0.9996); the transverse velocity *v* ∗ is the hardest component.

### 3.2 Supervised-data fraction and architecture

The supervised-data fraction, not the architecture, is the first-order driver of accuracy (Figure 10). At *df* ≥ 0.10 the backbones differ by one to two *R*^2^ points (Table 4), while at *df* ≤ 0.05 the spread widens sharply; across *df, R*^2^(*u*) climbs from ≈ 0 under pure physics (*df* = 0) to ≈ 0.95 by *df* = 0.01 and saturates near 0.97–0.99 by *df* = 0.20, the largest single gain falling between *df* = 0.001 and 0.01. For three of the four backbones, doubling the data from *df* = 0.10 to 0.20 raises *R*^2^(*u*) by only 0.001 to 0.015 (Table 4), so the sweep was not extended beyond 0.20. The Multi-Scale Fourier sweep spans *df* = 0.005 to 0.20 under both enforcement modes and shows the same shape, though the two modes separate in the scarce-data regime (at unequal budgets; Table S1): *R*^2^(*u*) = 0.43 (hard) and 0.70 (soft) at *df* = 0.005, 0.86 to 0.95 at *df* = 0.01, and 0.96 to 0.98 from *df* = 0.05 upward, where the two coincide. Low-data robustness is likewise not the property of one architecture: the MLP with soft enforcement retains *R*^2^(*u*) = 0.953 at *df* = 0.01 and 0.699 at 0.001 without collapsing, while SIREN reaches 0.938 (hard) or 0.893 (soft) at *df* = 0.01 and the Modified Fourier network degrades below *df* = 0.10. Training budget is a bounded confounding factor: extending the Multi-Scale Fourier runs from 350,000 to 1,000,000 steps changes *R*^2^(*u*) by at most 0.021 and occasionally lowers it, so ≈ 300,000 steps is almost converged for *df* ≥ 0.01 and the matched comparison is not budget limited.

**Figure 10.**
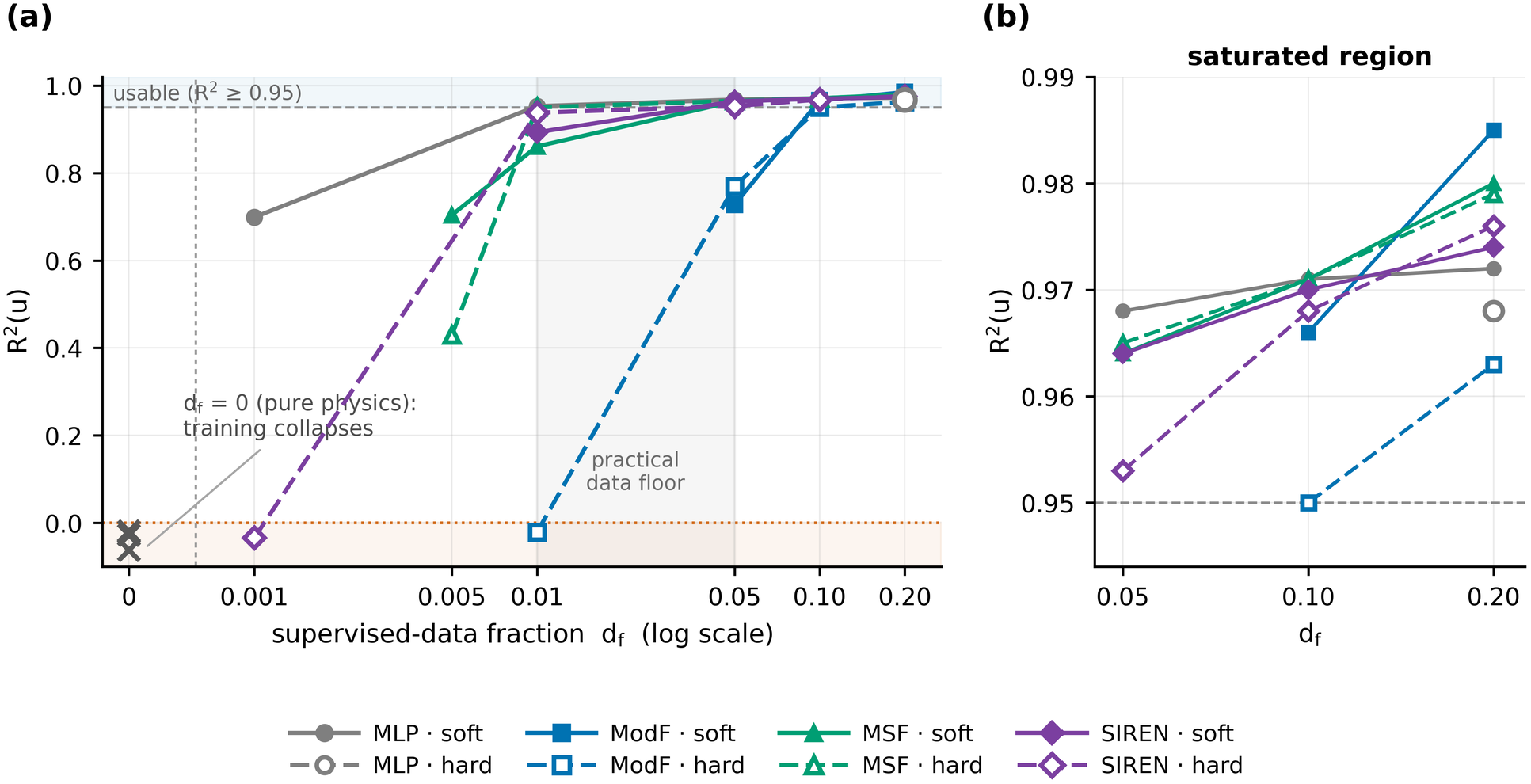
Validator *R*^2^(*u*) versus supervised-data fraction *df* for every architecture and inlet boundary condition. (a) Full sweep. The *df* = 0 vertical line is a pseudo-axis position beyond the dashed break; curves rise from *R*^2^ ≤ 0 under pure physics to ≈ 0.95 by *df* = 0.01 for the best backbones and cluster in the usable band by *df* = 0.20, so df is the first-order driver. The vertical band marks the practical data floor. (b) The saturated region, *df* ≥ 0.05, is shown on an expanded axis; each curve is drawn only over the range where its *R*^2^(*u*) lies inside the window, so a run that starts below the window appears only once it rises into view. Soft enforcement is drawn with a solid line and a filled marker, hard enforcement with a dashed line and an open marker. The three × at df = 0 are the data-free runs of Table S1, and the vertical axis stops just below the lowest value recorded in the study, *R*^2^ = −0.062(the data-free Multi-Scale Fourier run at Re ≈ 12.5).

Across all architectures, accuracy is governed first and foremost by the supervised-data fraction (Table 4). Every configuration reaches *R*^2^(*u*) ≥ 0.96 at *df* = 0.20 and remains usable at *df* = 0.10, but a cliff appears between *df* = 0.10 and *df* = 0.05 for the Modified Fourier network, and a more severe one between *df* = 0.05 and *df* = 0.01. The practical floor for soft-constrained training on this 46-outlet geometry is therefore *df* ≈ 0.05–0.10, roughly 5,000 supervised points, or one per 0.5 mm^3^ of lumen, below which default training does not reliably reach the correct flow.

### 3.3 Failure mode below the data floor

Boundary-condition enforcement matters mainly in the scarce-data regime, which is the regime this work targets. At *df* ≥ 0.10 soft and hard inlet enforcement are indistinguishable: SIREN at *df* = 0.20 reaches *R*^2^(*u*) = 0.974 soft versus 0.976 hard (at unequal budgets; Figures S2 and S3), and the Multi-Scale Fourier network 0.979 versus 0.978 at a matched 300,000 steps. The MLP behaves the same way (0.972 soft at 500,000 steps versus 0.968 hard at 300,000 steps), confirming that architectural inlet enforcement is not required for that backbone. At *df* = 0.01 the difference appears for the Fourier-family and SIREN backbones, whose soft inlet under-shoots 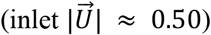 hard enforcement raises SIREN from 0.893 to 0.938 and the Multi-Scale Fourier network from 0.861 to 0.930–0.950, locking the inlet magnitude near 1.05 (Figures S4 and S5; the budgets differ, see Table S1). Hard enforcement is a useful safeguard at low data but is not sufficient alone: applied to the Modified Fourier backbone at *df* = 0.01 it did not recover within a 23,000-step budget, because removing the inlet-trivial mode does not prevent the bulk-trivial mode. Spatial fields and per-field accuracy for the remaining configurations are given in Figures S8 to S14.

The failure mode below the data floor is a specific, diagnosable attractor. The constant state 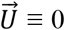 with uniform pressure satisfies continuity (∇ · 0 = 0), momentum (all convective and viscous terms vanish), and no-slip exactly:

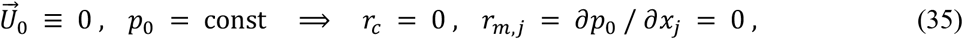

so only the inlet and data terms distinguish the trivial state from the true flow. When those terms are weak, training can settle near the trivial state, a known attractor of PINN training [17]. Empirically, collapsed runs show predicted velocity standard deviations of only ≈ 5–7% of the true field, an inlet velocity magnitude near 0.01 instead of 1.0, and the predicted mean velocity falls toward zero through the vessel bulk (Figure 11). A purely data-free configuration (*df* = 0) with hard-BC SIREN did not escape this attractor (*R*^2^ = −0.027, −0.029, −0.625, −0.180 for *u, v, w, p* at 142,700 steps), and neither did the two other data-free runs of Table S1. Its aggregated training loss was lower than that of the working df = 0.01 run, but only that run carries a data term; on the same collocation points with the training weights, the physics-and-boundary loss of a collapsed network is far higher than that of a converged one (21 for SIREN at df = 0.001 against 0.61 at df = 0.20), so collapsed runs sit on a high-loss plateau that the optimizer does not leave, not at a lower minimum.

**Figure 11.**
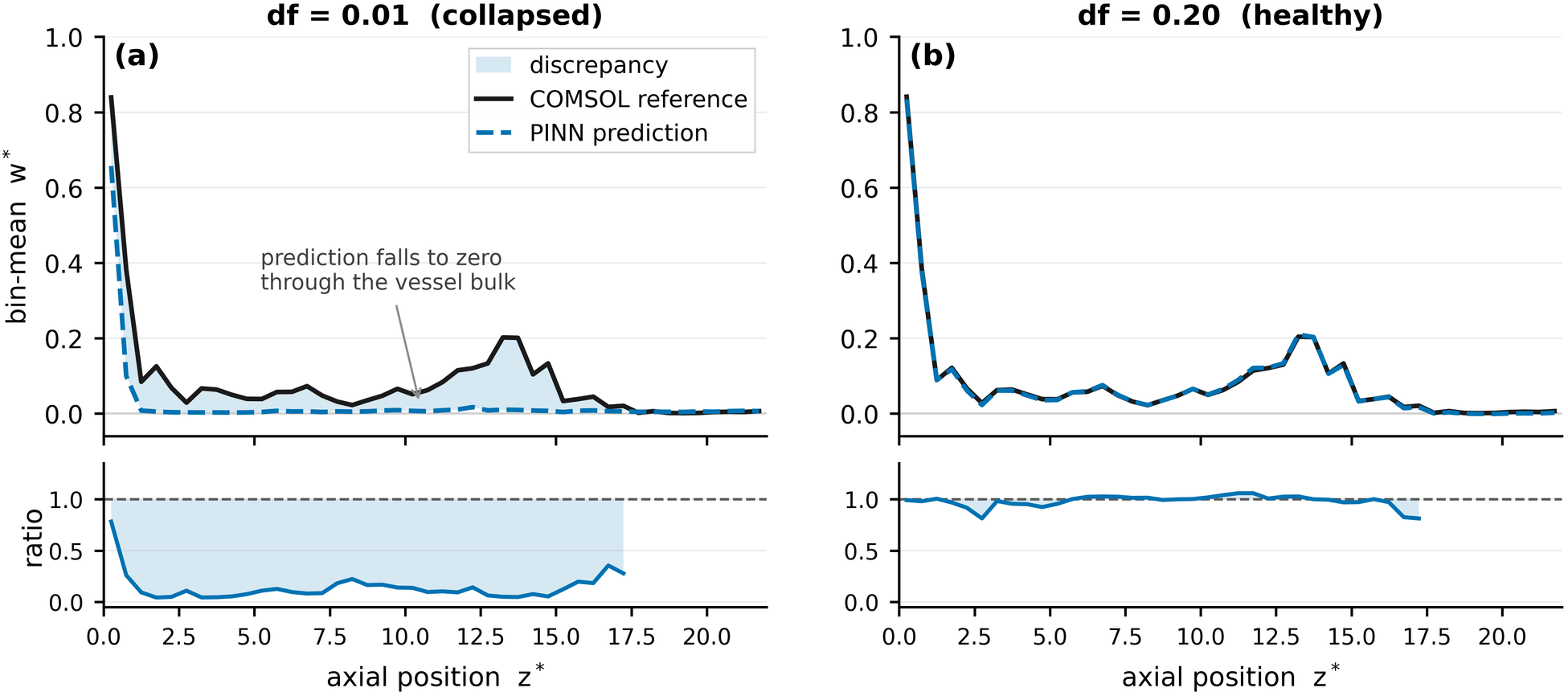
The trivial-solution attractor, on Modified Fourier runs. Bin-mean streamwise velocity against axial position at (a) *df* = 0.01, where the prediction falls to zero through the vessel bulk while satisfying the interior PDE residual, and (b) *df* = 0.20, where it tracks the reference. Shading marks the discrepancy; the strip beneath gives the predicted-to-reference ratio, undefined where the reference vanishes. Median ratio 0.11 in (a) against 1.00 in (b).

### 3.4 Absorbed-dose comparison

To test whether the surrogate’s flow field is clinically actionable we propagated it through a Y-90 microsphere transport and dosimetry pipeline (1.14 million spheres carrying 2.85 GBq, partitioned by the flow split between outlets and grouped into Couinaud segments V–VIII plus a tumor territory), using the reference *df* = 0.20 Multi-Scale Fourier checkpoint. The per-outlet flow split matches CFD with a Pearson correlation of 0.999 and an aggregate *L*^1^ discrepancy of 4.4 percentage points. The total outflow agrees to 2.7%. Because *L*^1^ sums unsigned errors over 46 outlets its maximum is 200%, so 4.4 points is a mean per-outlet error below 0.1 points. Propagated through the dose calculation (Table 5) the endpoints agree closely: perfused volume 149 versus 152 mL (−2.0%), mean perfused dose 896.1 versus 880.5 Gy (+1.8%), and every isodose volume from *V*_50_ to *V*_400_ within 1 mL. Tumor activity agrees to 5.6% and the four segment activities to within 3.2%. In both runs the deposited energy per unit organ mass (138.0 Gy) matches the MIRD estimate (137.8 Gy), a closure check on the kernel normalization. Two outlets returned small negative fluxes totaling 0.004% of the outflow; the pipeline clips these to zero, so they received no microspheres. Figure 12 compares the spatial dose distributions from the two flow splits as maximum-intensity projections; the deposits coincide branch for branch, with no territory systematically over-or under-dosed, and Figure 13 summarizes the dose-volume histogram and the delivered activity per territory.

**Table 5.**
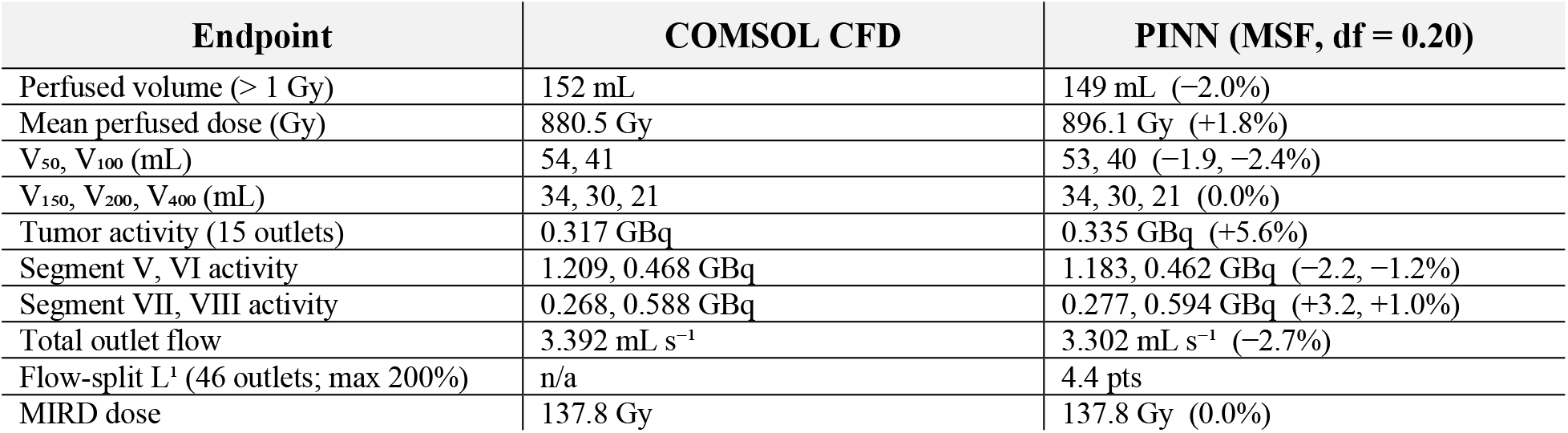
End-to-end Y-90 dosimetry, COMSOL CFD versus the PINN surrogate on the reference df = 0.20 Multi-Scale Fourier checkpoint (300,000 steps). Both calculations place 1,140,000 glass microspheres carrying 2.85 GBq and use the identical 1.0 mm dose grid and Y-90 dose-point kernel, differing only in the flow split. Outlet counts per territory: tumor 15, segment V 6, VI 10, VII 2, VIII 13. Percentages are computed from unrounded values.

**Figure 12.**
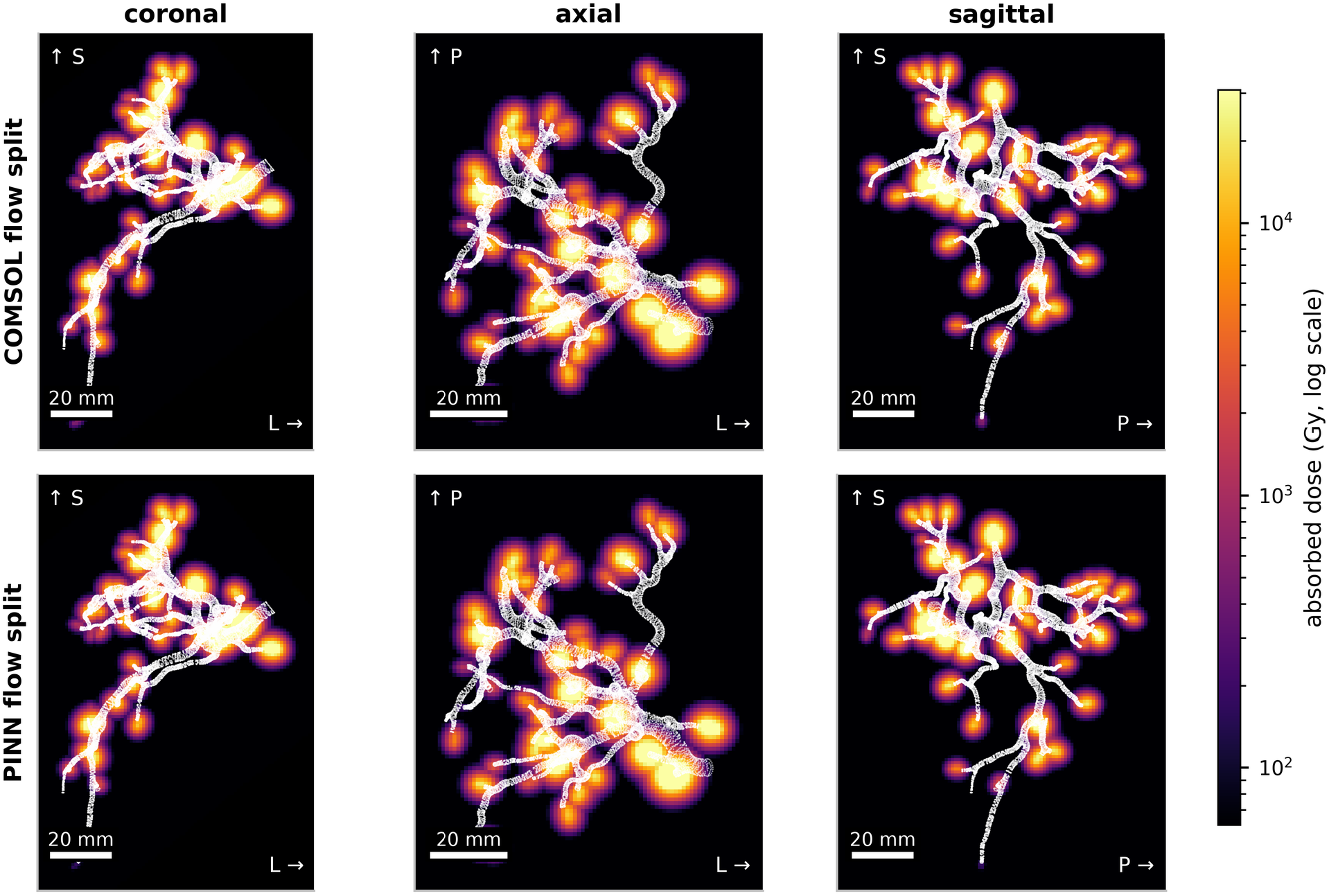
Coronal, axial and sagittal maximum-intensity projections of the Y-90 absorbed-dose distribution from the COMSOL flow split (top row) and the PINN flow split (bottom row), using the identical dose-point-kernel pipeline and a common logarithmic color scale, with the arterial tree overlaid in white. The dose volume is accumulated on the rotated solver grid and resampled into the CT frame of Figure 3, conserving total deposited energy to within 0.1%. The per-voxel maximum is a placement artifact, so only the dose-volume metrics of Figure 13 and Table 5 are quantitative. Scale bar 20 mm.

**Figure 13.**
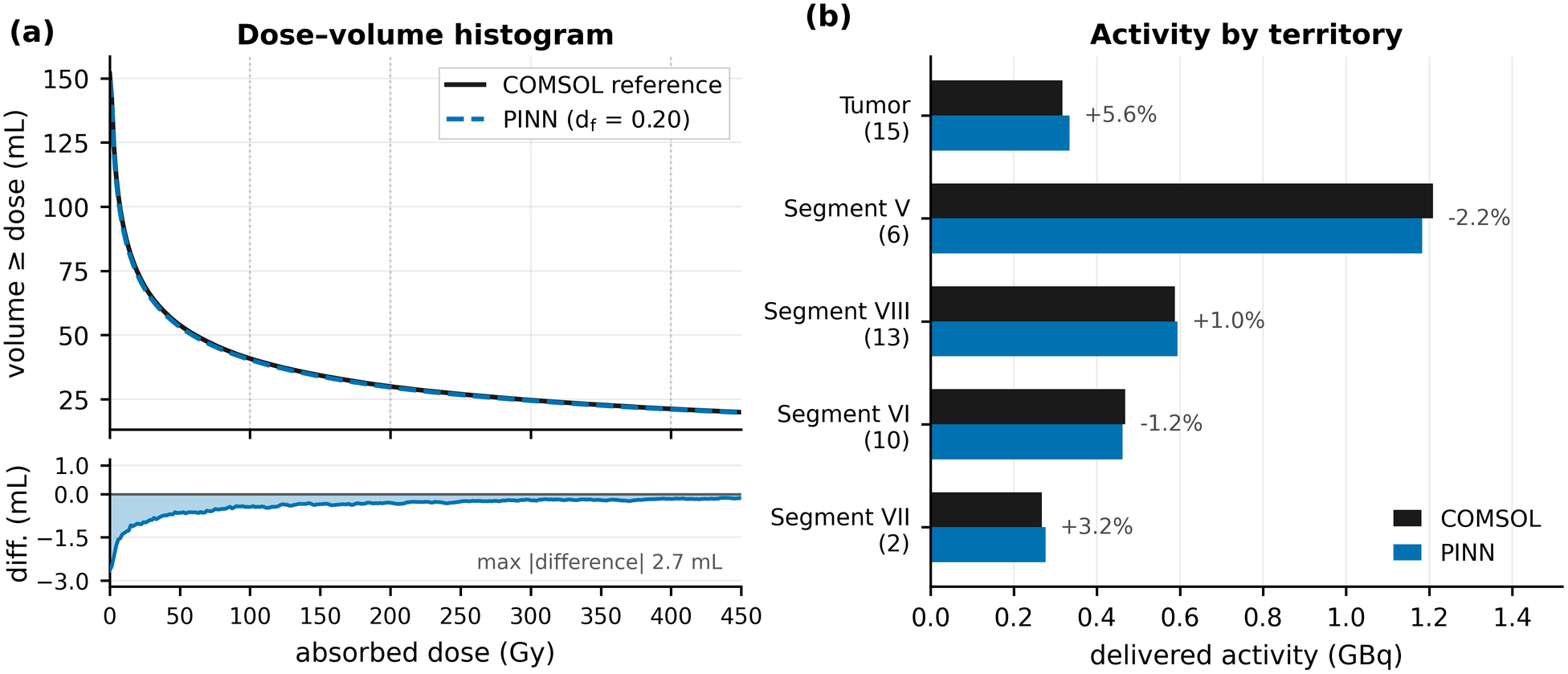
End-to-end agreement between the PINN surrogate (Multi-Scale Fourier, *df* = 0.20) and COMSOL. (a) Cumulative dose-volume histogram with the PINN-minus-COMSOL difference beneath. (b) Delivered activity per territory from 2.85 GBq, annotated with the relative difference; feeding-outlet counts in parentheses. The territory totals behind both panels are listed in Table 5.

Per-outlet errors partly cancel when summed into a territory, which is why a 4.4-point *L*^1^ flow-split discrepancy still leaves the dose-volume endpoints of Table 5 within 2.4%. The cancellation is not uniform, however, and it does not scale with territory size or with the number of feeding outlets: territory activities agree to within 5.6%, the largest difference being for the tumor, fed by 15 outlets (+5.6%), while segment VII, fed by two outlets, is at +3.2%. This does not mean that more feeding outlets bring larger errors: segments VI and VIII, fed by 10 and 13 outlets, agree to within 1.2%, while 72% of the tumor difference comes from a single one of its 15 outlets. A territory’s error therefore reflects how well its main outlets are split rather than how many outlets feed it. On a low-data checkpoint (MLP, *df* = 0.001, *R*^2^(*u*) = 0.70) the same pipeline gave an *L*^1^ discrepancy of 58.8 points and a 59% throughput deficit, confirming that the flow split, not the dose model, is error-limiting.

## 4. Discussion

Our results indicate that the supervised-data fraction is the primary design lever for this 3D hepatic artery at this Reynolds number (*Re* = 250.7) rather than the coordinate-network architecture. At *df* ≥ 0.10 all four architectures and both enforcement modes agree to within one to two *R*^2^ points. At a step-matched budget of *df* = 0.20 (Table 3, Figure 7) the difference between Multi-Scale Fourier, SIREN, and MLP (0.978, 0.976, 0.968, respectively) is much smaller than a single step along the *df* axis. The Modified Fourier network reached 0.985 only at 1.8 times that budget, and 0.963 inside it. Because tripling the budget changes *R*^2^(*u*) by up to 0.021 in either direction, the gaps of about 0.01 between backbones are within the effect of training length and too small to matter. The weight-normalized MLP remains a strong baseline, degrading under data scarcity but recovering the field from 1% data without architectural inlet enforcement. The backbone mattered only in the scarce-data regime.

The most data-efficient backbones recovered an accurate velocity field from as little as 1% of a converged CFD field (MLP with soft enforcement, *R*^2^(*u*) = 0.953 at *df* = 0.01; Multi-Scale Fourier with hard enforcement, 0.950), while others needed more data. All four saturate by *df* = 0.05 to 0.10; beyond that, R^2^(u) rises by at most 0.02. At *df* = 0, training collapsed toward the trivial state. This data floor is consistent with the data-anchored designs of other vascular PINNs [11], [14] and the convection-and stiffness-related optimization difficulties documented more generally [16]. Pressure is the least data-limited field because the outlet condition fixes its scale; velocity, which drives particle transport, is the data-limited quantity. We therefore suggest budgeting 5–10% supervised CFD points for a backbone-independent margin, and no less than 1% even for the most efficient network.

Even in the reference model, the residual error is not uniform. The largest absolute velocity errors cluster at the proximal bifurcation, roughly two inlet diameters downstream of the inlet, where the flow splits over a short length scale and spatial gradients are several times the domain average. Within the first eight inlet diameters along the vessel, the mean error is 2.4% to 3.0% of the field range against 0.64% over the whole domain, and this region, although only 2.8% of the validation points, holds 36% of the largest 1% of errors. Distal branches, with smoother and slower flow, are more accurate than the global figure (0.53%). Because the flow split at each bifurcation depends on the velocity field near the branch entrances, this localization is the error most relevant to dosimetry and a natural target for spatially weighted residuals in future work [29].

Our results are consistent with, and extend, prior PINN hemodynamics studies. Kissas et al. recovered arterial pressure waveforms and Windkessel parameters from 4D-flow MRI in arterial networks [10]; Sarabian et al. combined sparse clinical measurements with reduced-order physics to improve cerebral hemodynamic predictions, and Garay et al. estimated three-element Windkessel parameters from synthetic phase-contrast MRI [11], [12]; Daneker et al. used transfer learning to track a synthetically evolving false lumen [13]. PINN surrogates have likewise been reported for morphology-conditioned 4D hemodynamics [14], pressure estimation in an idealized aortic arch [30], and idealized aneurysm flow [31], while Moser et al. found no significant gain from Fourier-feature encoding over a fully connected network for three-dimensional blood flow [21], and Talaat et al. found near-wall treatment and nondimensionalization critical for three-dimensional airway flow [32]; in our sweep, too, architecture is secondary to the data fraction. What this study adds is, to our knowledge, the first systematic sweep of architecture, inlet enforcement and supervised-data fraction on a realistic multi-outlet hepatic tree, and the explicit demonstration and diagnosis of the trivial-solution floor.

Fast inference would make individual patient exploration and intra-procedural re-planning more feasible once the network generalizes across catheter positions and boundary conditions, a prerequisite for closed-loop catheter optimization. The natural end-to-end validation is to integrate microsphere trajectories on the PINN-predicted field and compare the resulting activity distribution with post-treatment Y-90 PET/CT, following the CFD-dosimetry pipeline previously demonstrated for this application [6]. Operator-learning surrogates that generalize across patient geometries [9] will be developed in the future.

The main limitations of this first report of a PINN surrogate for Y-90 radioembolization include using a single patient geometry, a single inlet flow rate, a single uniform zero-pressure outlet model, steady flow, rigid walls, Newtonian blood, and a single random seed. Because the flow split and hence the dose are sensitive to downstream resistance [2], replacing the uniform *p* = 0 condition with a three-element Windkessel boundary condition at each outlet, which PINNs can also infer from the data [12], is the most important next step. The data-efficiency floor was also characterized without adaptive loss balancing [26] or Bayesian uncertainty quantification [33], either of which could lower it and attach confidence intervals to predictions. Further priorities are an inlet-flow and Reynolds sweep; pulsatile flow with a cardiac waveform; mechanisms to make the data-free regime tractable, such as spatially concentrated continuity weighting, pseudo-time continuation [34] or per-outlet flux constraints; multi-patient operator learning [9]; and spatially weighted residuals targeting the proximal-bifurcation error [29]. Domain-decomposition PINNs [35] and the training stack developed for three-dimensional turbulence [36] offer routes to higher Reynolds numbers and larger trees.

## 5. Conclusion

We presented a patient-specific PINN model for hepatic arterial hemodynamics in close agreement with the reference CFD solution (*R*^2^(*u*) > 0.97, *R*^2^ ≥ 0.96 for every velocity component and ≈ 1.0 for pressure at 20% supervision) and full-field inference in seconds. Matched on both data fraction and training budget, the architectures are closely comparable, showing that accuracy is governed primarily by the supervised-data fraction. Using our Y-90 dosimetry pipeline with 20% CFD data, the PINN reproduced the CFD perfused volume, mean dose and isodose volumes within 2.4%, the segment activities within 3.2% and the tumor activity within 5.6%. These results give data budget, boundary condition and architecture guidelines for PINN surrogates in Y-90 radioembolization planning.

## Acknowledgments

We thank UC Davis Health for the patient geometry. NVIDIA PhysicsNeMo and PhysicsNeMo Sym (v2) were used for all training. Claude (Anthropic) was used for code-development support and language editing; the authors take full responsibility for all scientific content.

## Data and code availability

The training scripts, Hydra configurations, flow-split extraction and the Y-90 dosimetry pipeline needed to reproduce every number reported here are available from the corresponding author on reasonable request.

## Author contributions

E.T. designed and ran the PINNs study, implemented the dosimetry pipeline and wrote the manuscript. N.K.P.S. produced the COMSOL reference solutions. E.R. supervised the work and revised the manuscript. All authors approved the final version.

