## Supplementary Material (Notes S1, Figures S1-S14, Table S1) for "A Physics-Informed Neural Network Surrogate for Patient-Specific Hepatic Arterial Hemodynamics in Yttrium-90 Radioembolization: Network Architecture, Boundary-Condition Enforcement, and Data Efficiency"

This document contains the extended methods of Supplementary Note S1 (Eqs. S1–S24), the Y-90 dose-point kernel used for every dose map (Figure S1), the per-architecture field and accuracy figures referenced in the main text (Figures S2–S14) and the complete run-by-run results table (Table S1). All runs share the same patient geometry, and all but one data-free run share the same flow (Re ≈ 251); df denotes the supervised-data fraction and $R^2$ is the validator value $1 - (\text{relative } L^2)^2$ at the final training step.

### Supplementary Note S1. Extended methods

This note collects the formal definitions that support Section 2 of the main text: the four coordinate-network backbones, the inlet profile and boundary-condition implementation, the optimizer, the identity relating the two accuracy metrics, and the microsphere-placement and dose-accumulation geometry. Symbols follow the main text: an asterisk marks a nondimensional quantity, and $q_{ref}$ denotes the COMSOL reference field.

#### S1.1 Coordinate-network architectures

The baseline multilayer perceptron maps coordinates to fields through six SiLU-activated hidden layers of width 256, with every linear layer reparameterized by weight normalization:

$$h^0 = \sigma(W^0 x^* + b^0), h^{\ell+1} = \sigma\left(W^{\ell+1} h^\ell + b^{\ell+1}\right), (u^*, v^*, w^*, p^*)^T = W^o h^L + b^o, \tag{S1}$$

$$\sigma(x) = x \cdot (1 + e^{-x})^{-1}, \tag{S2}$$

$$W = g \cdot v / \|v\|. \tag{S3}$$

The Modified Fourier network first lifts the input through an axis-aligned Fourier-feature embedding with 18 frequencies F per axis, then mixes two parallel encoders through per-layer multiplicative gates, where $\odot$ denotes the elementwise (Hadamard) product:

$$\gamma(x^*) = [x^*, sin(2\pi F x^*), cos(2\pi F x^*)]^T, \tag{S4}$$

$$U = \sigma(W_U \gamma(x^*) + b_{UU}), V = \sigma(W_V \gamma(x^*) + b_V), \tag{S5}$$

$$Z^\ell = \sigma\left(W^\ell H^\ell + b^\ell\right), \tag{S6}$$

$$H^{\ell+1} = \left(1 - Z^\ell\right) \odot U + Z^\ell \odot V. \tag{S7}$$

The Multi-Scale Fourier network applies three independent axis-aligned Fourier encodings, one per frequency band, to the same coordinates in parallel. A single trunk with shared weights processes each encoding and the band outputs are concatenated before one final linear layer:

$$\gamma_{\mathrm{i}}(x^*) = [\, x^*,\ sin(2\pi F_{\mathrm{i}} x^*),\ cos(2\pi F_{\mathrm{i}} x^*)\,]^{\mathrm{T}},\ i = 1, 2, 3, \tag{S8}$$

$$h_{\mathrm{i}} = f\theta(\gamma_{\mathrm{i}}(x^*)),\ \ \Phi(x^*) = W\,[\, h_1\,;\ h_2\,;\ h_3\,] + b. \tag{S9}$$

The sinusoidal-representation network composes sine-activated layers under the frequency-scaled initialization of Sitzmann et al., which keeps the pre-activation distribution invariant with depth:

$$\Phi(x^*) = W^L(\varphi_{L-1} \circ \cdots \circ \varphi_0)(x^*) + b_n^L,\ \varphi_\ell(z) = \sin\left(\omega_\ell(W_\ell z + b_\ell)\right), \tag{S10}$$

with first-layer frequency $\omega_0 = 30$ and hidden frequency $\omega = 10$, and the variance-preserving initialization

$$W_\ell \sim U\left(-\sqrt{(6/n_\ell)}/\omega,\ +\sqrt{(6/n_\ell)}/\omega\right). \tag{S11}$$

#### S1.2 Inlet profile and boundary conditions

The prescribed inlet profile is a power law of exponent n, auto-fitted to the near-inlet reference data and normalized so that its area average equals the bulk velocity, with R the inlet radius and r the in-plane radius:

$$u_{in}^*(r) = u_{peak}^*[1 - (r/R)^n], \tag{S12}$$

$$u^*_{peak} = \big((n+2)/n\big)U^*_m. \tag{S13}$$

Global mass conservation and the absence of swirl are imposed as an integral flux constraint and two tangential conditions, with the two tangent vectors spanning the inlet plane:

$$\int_{\Gamma_{in}} \big(\vec{U}^* \cdot n\big) dS = Q_{in}, \tag{S14}$$

$$\vec{U}^* \cdot t_1 = 0,\ \vec{U}^* \cdot t_2 = 0. \tag{S15}$$

Under hard inlet enforcement the geometry is rotated so that the inlet lies at z = 0 and the raw network outputs are blended toward the prescribed profile by a smooth gate of width δ:

$$\varphi(z) = \tanh(z/\delta), \tag{S16}$$

$$u^* = \varphi u^{*\prime}, v^* = \varphi v^{*\prime}, w^* = (1-\varphi) w^*_{in} + \varphi w^{*\prime}, \tag{S17}$$

$$w^*_{in}(x,y) = u^*_{peak} \cdot max(0,1-(r/R)^n), r = \sqrt{(x^2+y^2)}. \tag{S18}$$

The gate vanishes at the inlet plane, so the prescribed profile is recovered exactly, and saturates to unity in the bulk, where the network is unconstrained.

#### S1.3 Loss terms

The loss terms are defined term by term in Section 2.5 of the main text (Eqs. 15–22).

#### S1.4 Optimizer

Parameters are updated with Adam, whose bias-corrected first- and second-moment estimates give the step, under an exponentially decaying learning-rate schedule with a fixed decay interval:

$$\theta_{s+1} = \theta_s - \eta_s \frac{\hat{m}_s}{\sqrt{\hat{v}_s} + \varepsilon}, \tag{S19}$$

$$\eta_s = \eta_0 \cdot \gamma^{s/s_d}, \tag{S20}$$

where $\hat{m}_s = \frac{m_s}{1-\beta_1{}^s}$ and $\hat{v}_s = \frac{v_s}{1-\beta_2{}^s}$ are the bias-corrected moment estimates ($\beta_1 = 0.9$, $\beta_2 = 0.999$, $\varepsilon = 10^{-8}$), $\gamma = 0.95$ is the decay rate and $s_d$ = 20,000 steps the decay interval.

#### S1.5 Relation between the two accuracy metrics

The coefficient of determination and the relative $L^2$ error are equivalent up to a field-dependent constant that depends only on the ground truth:

$$\kappa(q) = \|q^{\mathrm{true}}\|^2 / \|q^{\mathrm{true}} - \bar{q}^{\mathrm{true}}\|^2, R^2(q) = 1 - \kappa(q)[L_{\mathrm{rel}}\, e^2(q)]^2. \tag{S21}$$

#### S1.6 Microsphere placement and dose accumulation

The spheres assigned to outlet k are treated as point sources of activity and placed uniformly at random in a cylinder of the cap-equivalent radius $R_k = \sqrt{(A_k/\pi)}$ and fixed length ℓ = 4 mm, which starts at the cap centroid $c_k$ and extends downstream along the outlet normal $n_k$; the sphere size does not enter:

$$p = c_k + \ell\xi_1 n_k + R_k\sqrt{\xi_2}(cos\,\theta\; e_1 + sin\,\theta\; e_2), \quad \theta = 2\pi\xi_3 \qquad \text{(S22)}$$

where $\xi_1$, $\xi_2$ and $\xi_3$ are independent random numbers uniform on [0, 1) and $e_1$, $e_2$ are orthonormal vectors in the cap plane. Each sphere is binned into the 1 mm voxel that contains it, so that each voxel carries an activity proportional to its sphere count:

$$a(x) \;=\; n_{sph}(x) \;\cdot\; a_s. \qquad \text{(S23)}$$

Because the dose-point kernel (Figure S1) integrates to the mean β energy per decay, the total deposited energy is conserved, which provides a built-in consistency check on the whole chain:

$$\rho V_{\text{vox}} \textstyle\sum_x D(x) \;=\; \eta \bar{\text{E}}_\beta \textstyle\sum_x \tilde{\text{N}}(x) \;=\; \eta \bar{\text{E}}_\beta A_{\text{tot}}\tau. \qquad \text{(S24)}$$

where ρ is the tissue density, $V_{\text{vox}}$ the voxel volume, η the MeV-to-joule conversion and $\tau = T_{1/2}/\ln 2 \approx 92.4$ h the mean life of Y-90, so that $A_{\text{tot}}\tau$ is the total number of decays.

#### Supplementary Figures

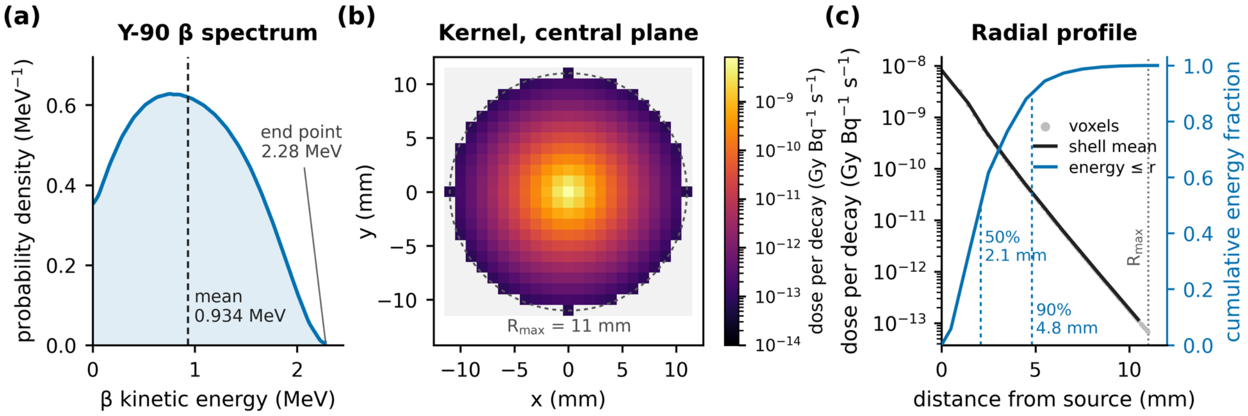


**Figure S1.** Y-90 source term and the dose-point kernel used for every dose map in this work. (a) Y-90 β spectrum as a probability density per MeV, replotted from Ref. [6] of the main text; its mean, 0.936 MeV, agrees with the mean β energy of 0.9337 MeV to which the kernel is normalized (dashed line, labeled 0.934 MeV), and it ends at 2.28 MeV. (b) Central plane of the three-dimensional kernel on the 1 mm dose grid: the analytic form of main-text Eq. (31) with α = 1.3 and λ = 1.1 mm, truncated at $R_{max}$ = 11 mm and normalized to deposit 0.9337 MeV per decay, shown as absorbed dose per decay for tissue of density 1.06 g/cm³ on a logarithmic scale. (c) The same kernel against distance from the source: individual voxels (grey), shell mean (black) and the cumulative fraction of the deposited energy (blue, right axis). Half of the energy is absorbed within 2.1 mm, 90% within 4.8 mm and none beyond 11 mm.

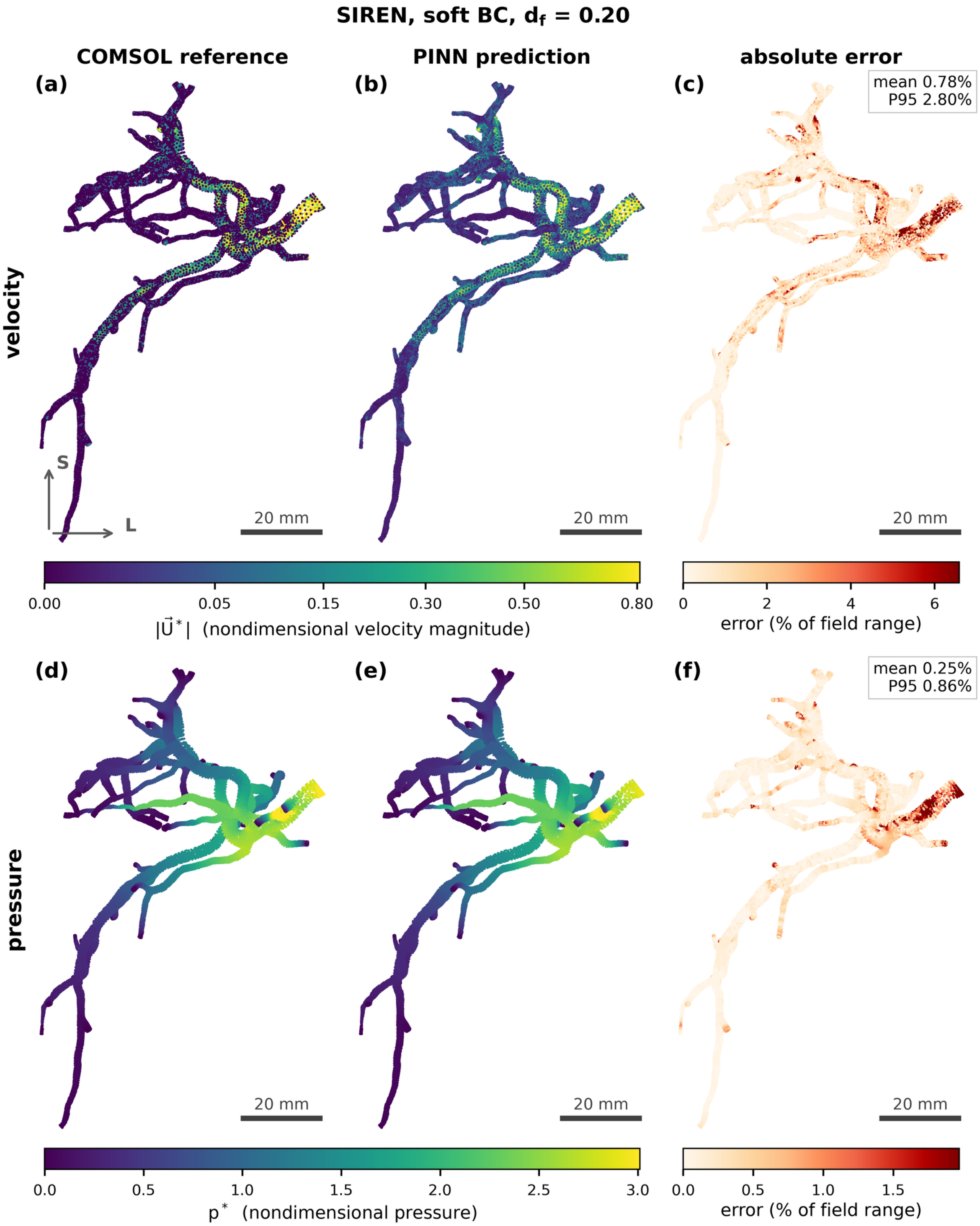


**Figure S2.** Spatial fields for the SIREN model with soft inlet enforcement at df = 0.20. Panels (a) to (c): velocity magnitude $|\vec{U}^*|$ for the COMSOL reference, the PINN prediction, and the absolute error as a percentage of the field range; panels (d) to (f): the same three views of the pressure p*. Anterior view in the CT frame, in the same orientation as main-text Figures 3 and 8: superior upward, the patient's right on the viewer's left. Scale bar 20 mm in every panel.

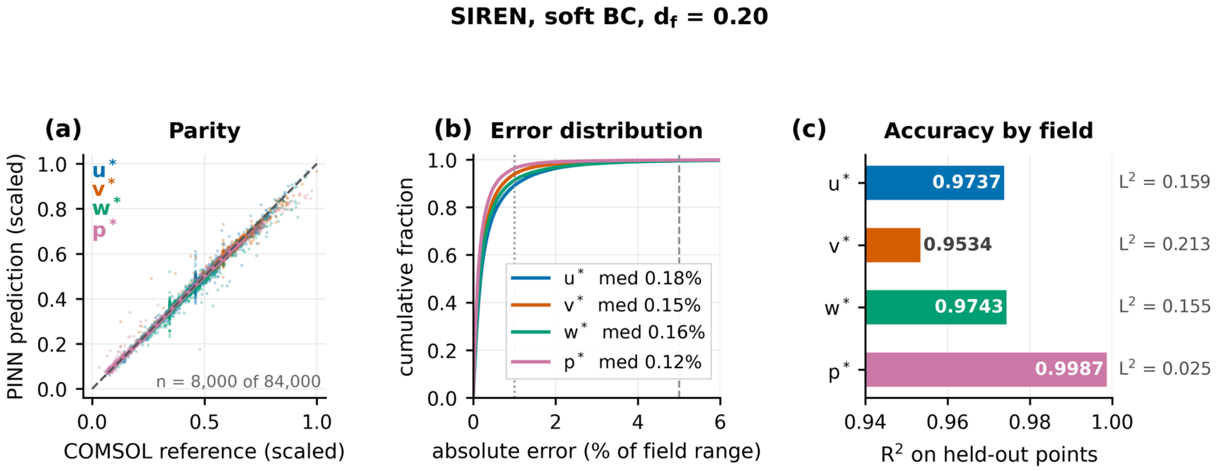


**Figure S3.** Accuracy of the soft-BC SIREN at df = 0.20: $R^2$ = 0.974 / 0.953 / 0.974 / 0.999 for u / v / w / p. Panels: (a) parity, with each field scaled to its own range; (b) cumulative distribution of the absolute error; (c) $R^2$ per field, with the relative $L^2$ printed alongside.

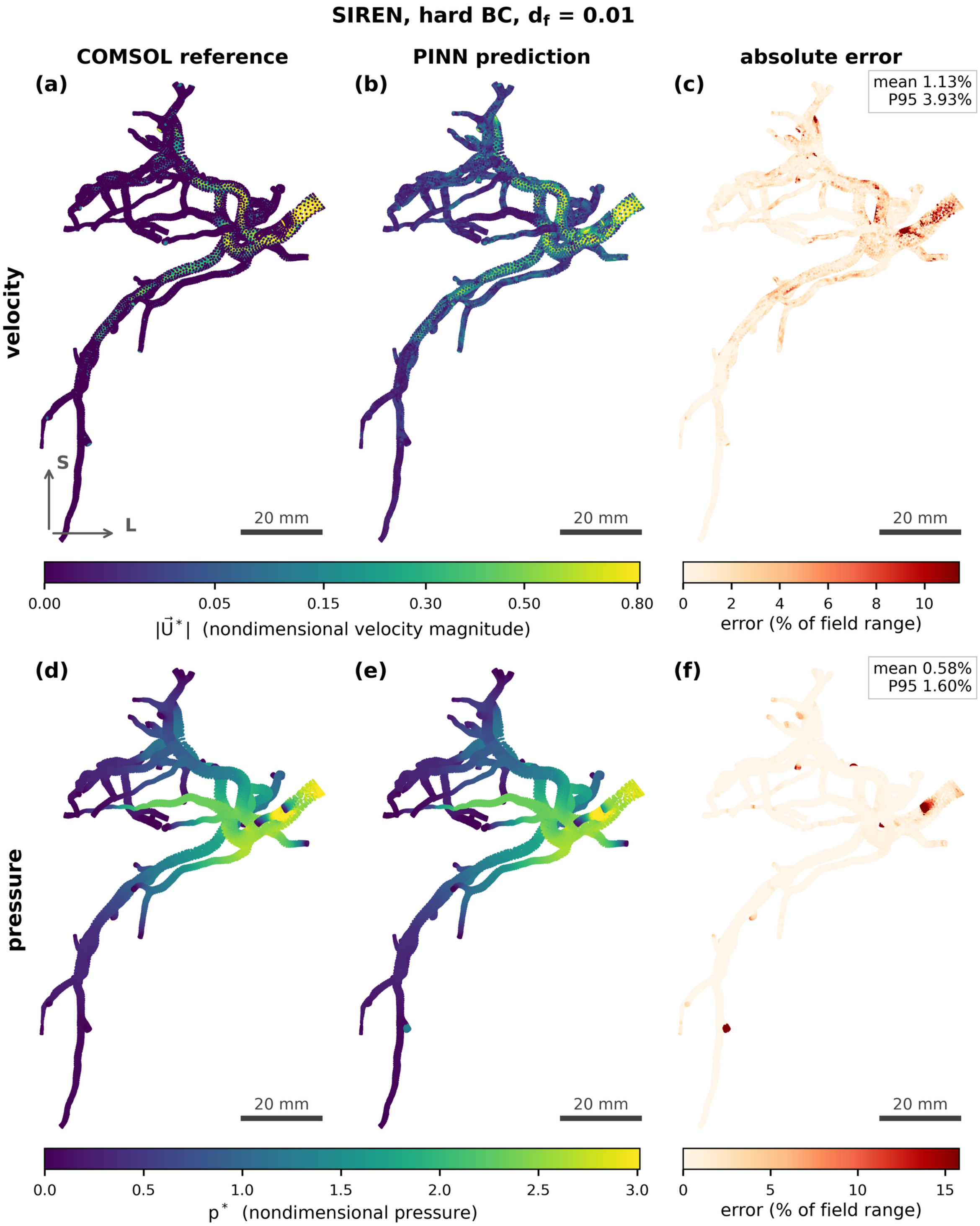


**Figure S4.** Spatial fields for the SIREN model with hard inlet enforcement at df = 0.01. Panels (a) to (c): velocity magnitude $|\vec{U}*|$ for the COMSOL reference, the PINN prediction, and the absolute error as a percentage of the field range; panels (d) to (f): the same three views of the pressure p*. Anterior view in the CT frame, in the same orientation as main-text Figures 3 and 8: superior upward, the patient's right on the viewer's left. Scale bar 20 mm in every panel.

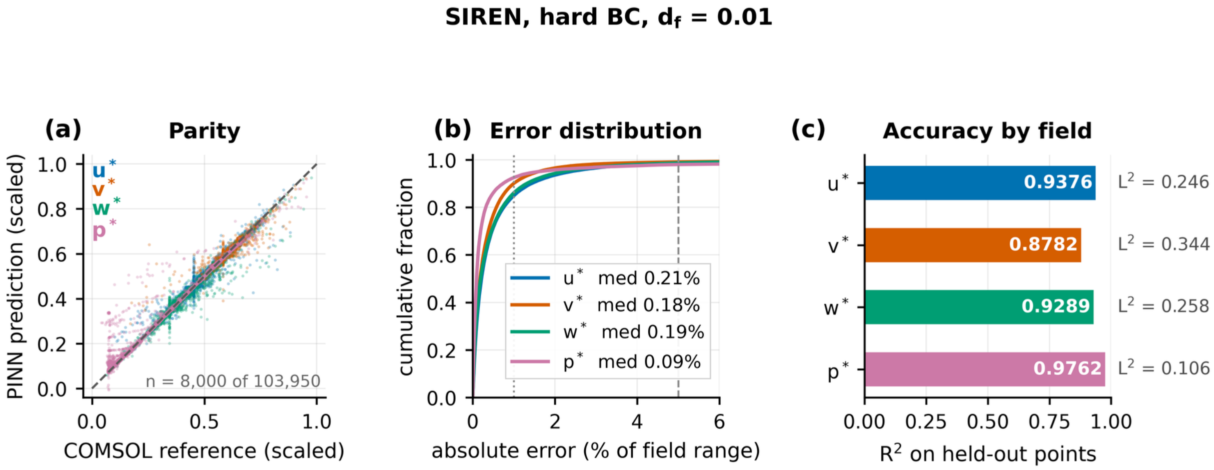


**Figure S5.** Accuracy of the hard-BC SIREN at df = 0.01: $R^2 = 0.938 / 0.878 / 0.929 / 0.976$. Hard inlet enforcement recovers the field from 1% supervised data. Panels: (a) parity, with each field scaled to its own range; (b) cumulative distribution of the absolute error; (c) $R^2$ per field, with the relative $L^2$ printed alongside.

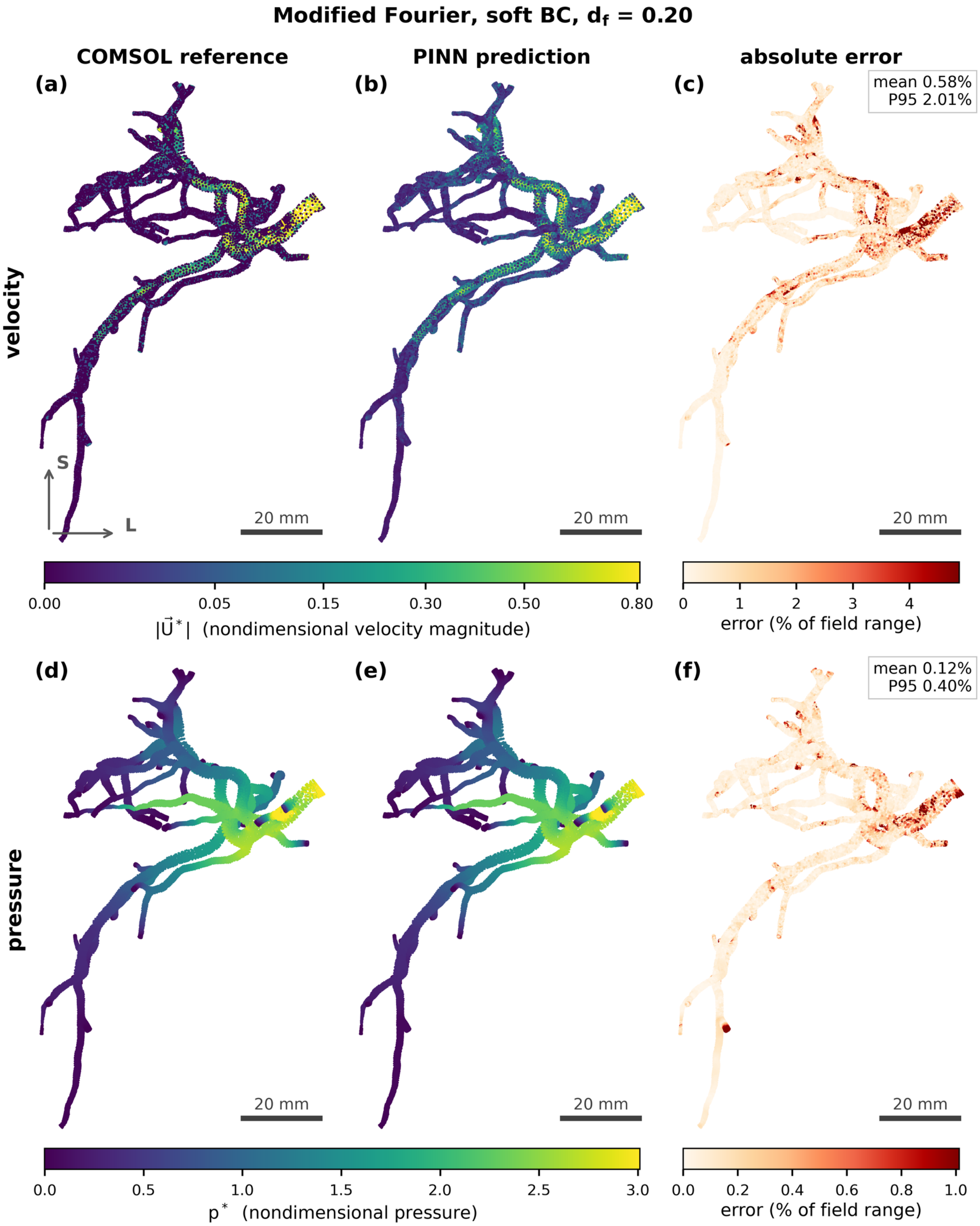


**Figure S6.** Spatial fields for the Modified Fourier model at df = 0.20, the highest-accuracy run in the study (536,000 steps). Panels (a) to (c): velocity magnitude $|\vec{U}*|$ for the COMSOL reference, the PINN prediction, and the absolute error as a percentage of the field range; panels (d) to (f): the same three views of the pressure p*. Anterior view in the CT frame, in the same orientation as main-text Figures 3 and 8: superior upward, the patient's right on the viewer's left. Scale bar 20 mm in every panel.

**Modified Fourier, soft BC, $d_f$ = 0.20**

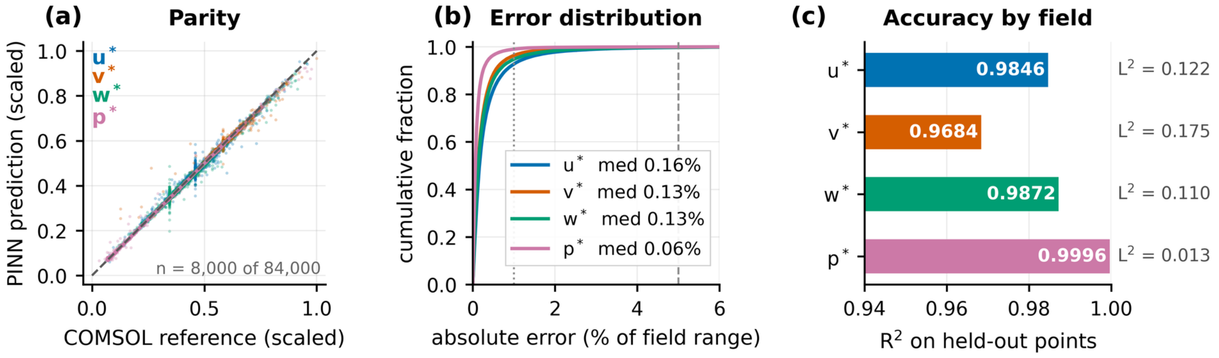


**Figure S7.** Accuracy of the Modified Fourier model at df = 0.20: $R^2$ = 0.985 / 0.968 / 0.987 / 1.000. Panels: (a) parity, with each field scaled to its own range; (b) cumulative distribution of the absolute error; (c) $R^2$ per field, with the relative $L^2$ printed alongside.

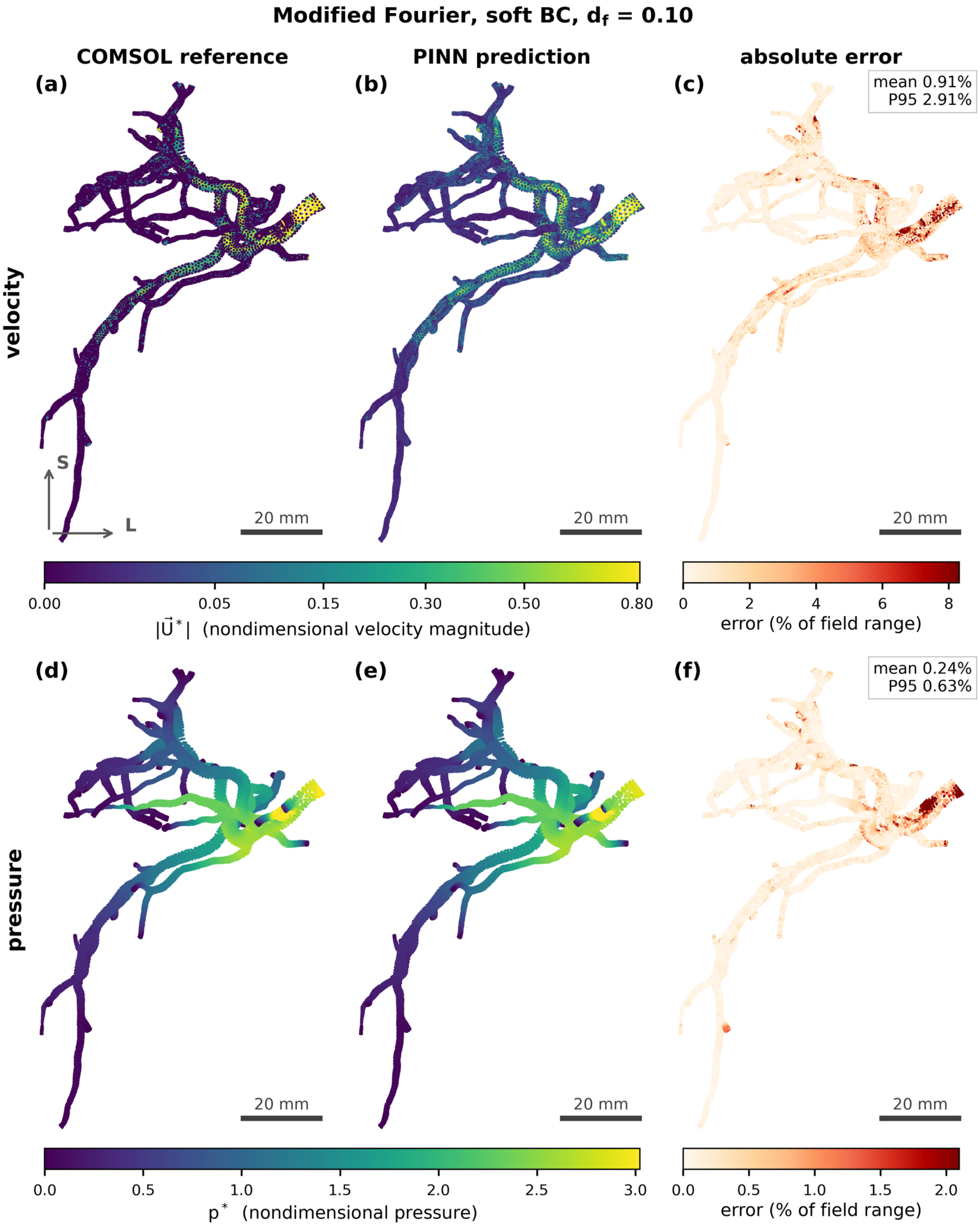


**Figure S8.** Spatial fields for the Modified Fourier model at df = 0.10, the lowest data fraction at which it retains strong accuracy. Panels (a) to (c): velocity magnitude $|\vec{U}*|$ for the COMSOL reference, the PINN prediction, and the absolute error as a percentage of the field range; panels (d) to (f): the same three views of the pressure p*. Anterior view in the CT frame, in the same orientation as main-text Figures 3 and 8: superior upward, the patient's right on the viewer's left. Scale bar 20 mm in every panel.

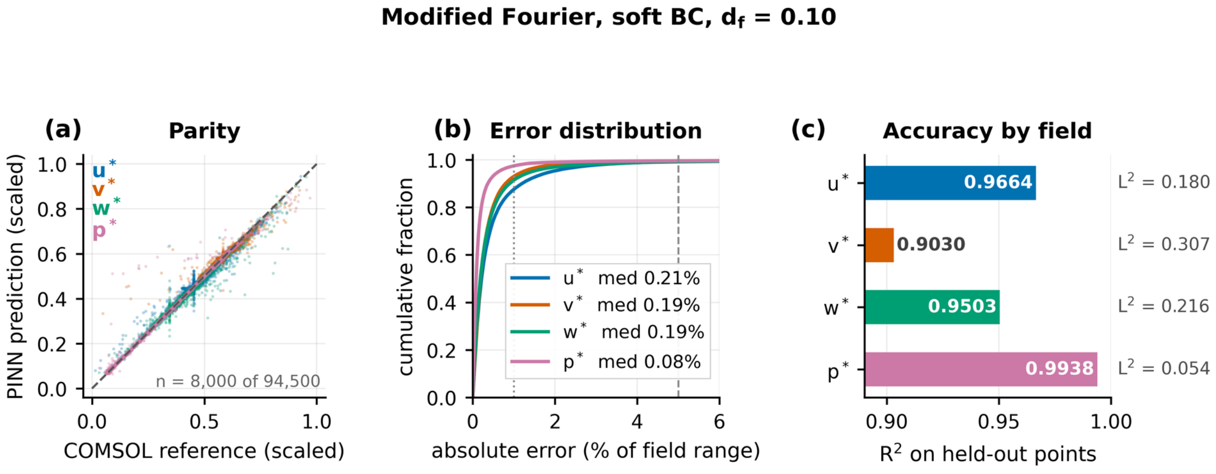


**Figure S9.** Accuracy of the Modified Fourier model at df = 0.10: $R^2$ = 0.966 / 0.903 / 0.950 / 0.994. Panels: (a) parity, with each field scaled to its own range; (b) cumulative distribution of the absolute error; (c) $R^2$ per field, with the relative $L^2$ printed alongside.

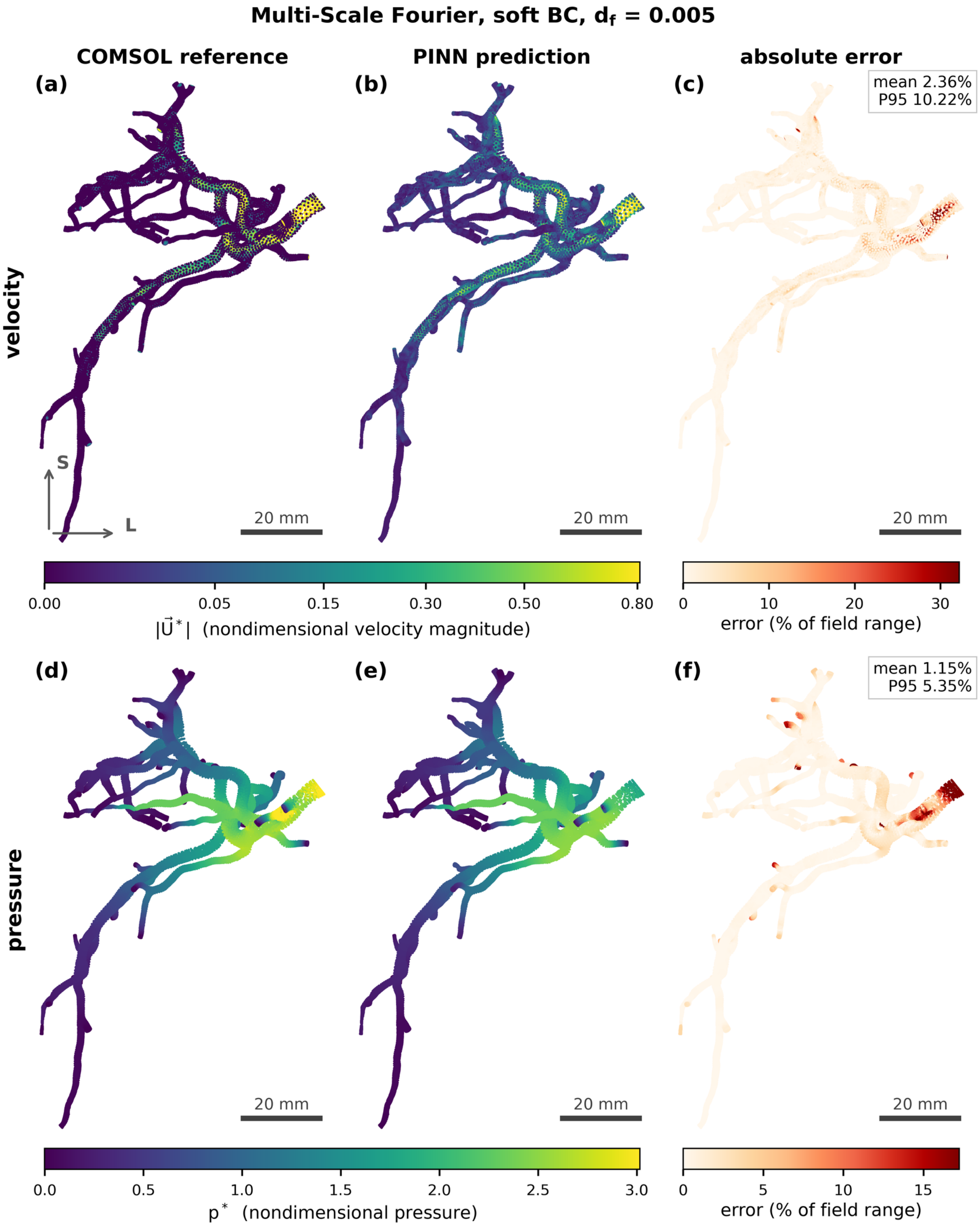


**Figure S10.** Spatial fields for the Multi-Scale Fourier model at df = 0.005 (2.05 M steps). Panels (a) to (c): velocity magnitude $|\vec{U}*|$ for the COMSOL reference, the PINN prediction, and the absolute error as a percentage of the field range; panels (d) to (f): the same three views of the pressure p*. Anterior view in the CT frame, in the same orientation as main-text Figures 3 and 8: superior upward, the patient's right on the viewer's left. Scale bar 20 mm in every panel.

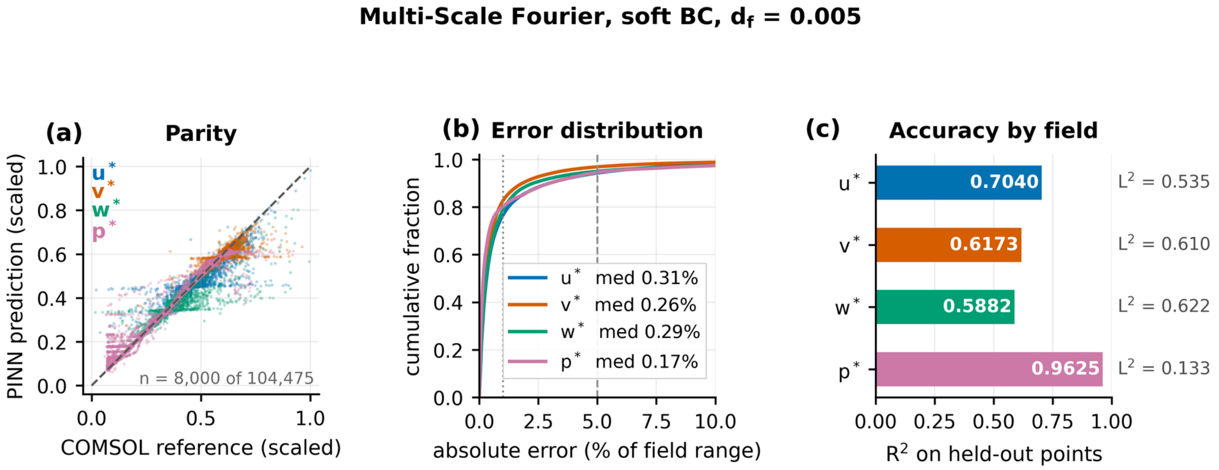


**Figure S11.** Accuracy of the Multi-Scale Fourier model at df = 0.005: $R^2$ = 0.704 / 0.617 / 0.588 / 0.963. Additional steps cannot compensate for missing data; note that panel (c) uses a full 0 to 1 axis. Panels: (a) parity, with each field scaled to its own range; (b) cumulative distribution of the absolute error; (c) $R^2$ per field, with the relative $L^2$ printed alongside.

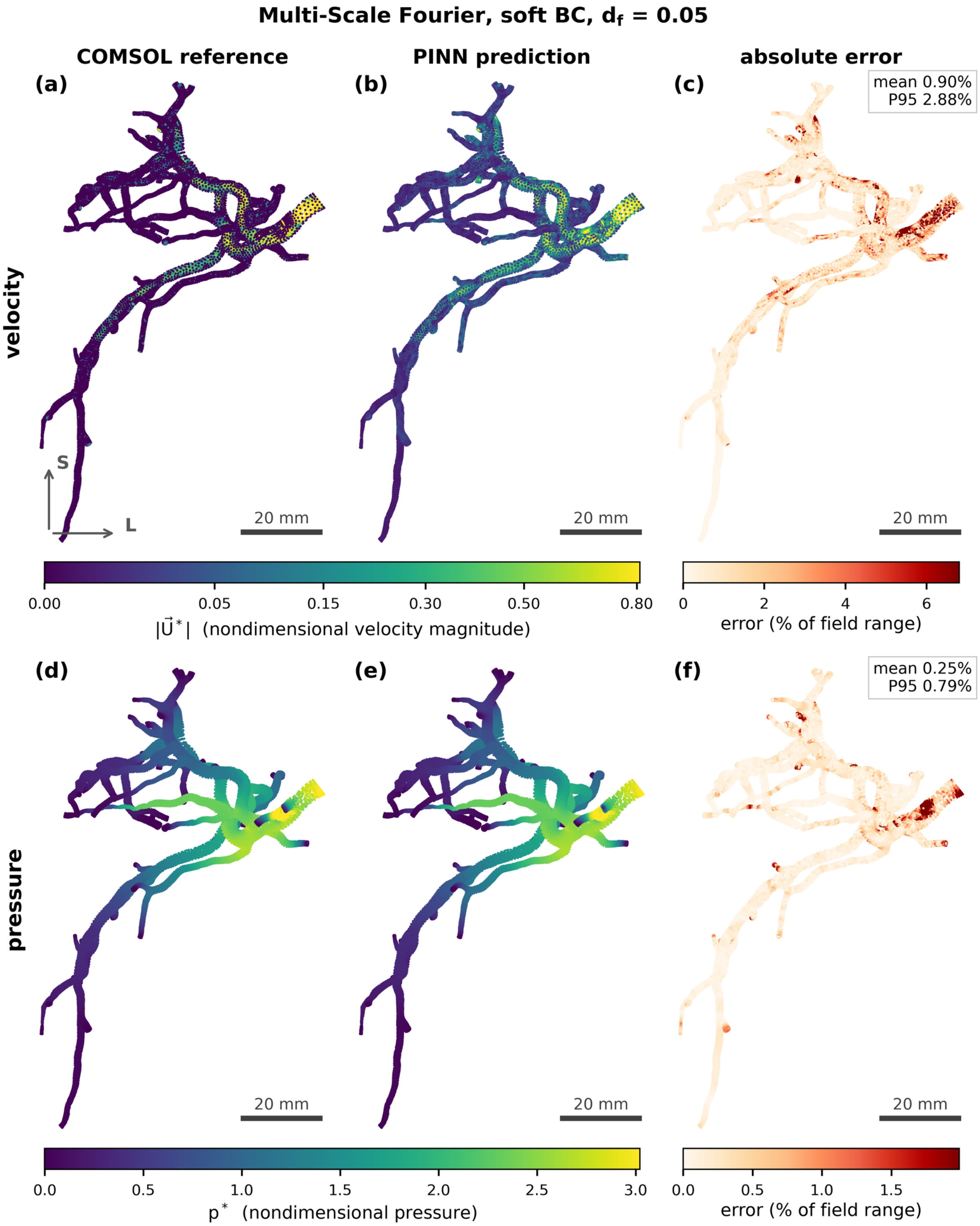


**Figure S12.** Spatial fields for the Multi-Scale Fourier model at df = 0.05. Panels (a) to (c): velocity magnitude $|\vec{U}*|$ for the COMSOL reference, the PINN prediction, and the absolute error as a percentage of the field range; panels (d) to (f): the same three views of the pressure p*. Anterior view in the CT frame, in the same orientation as main-text Figures 3 and 8: superior upward, the patient's right on the viewer's left. Scale bar 20 mm in every panel.

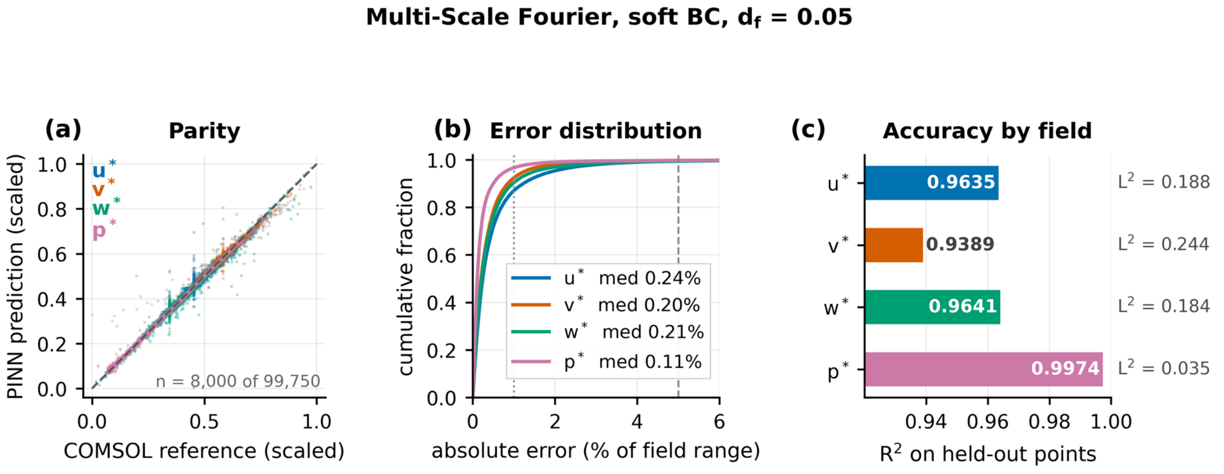


**Figure S13.** Accuracy of the Multi-Scale Fourier model at df = 0.05: $R^2$ = 0.964 / 0.939 / 0.964 / 0.997. Panels: (a) parity, with each field scaled to its own range; (b) cumulative distribution of the absolute error; (c) $R^2$ per field, with the relative $L^2$ printed alongside.

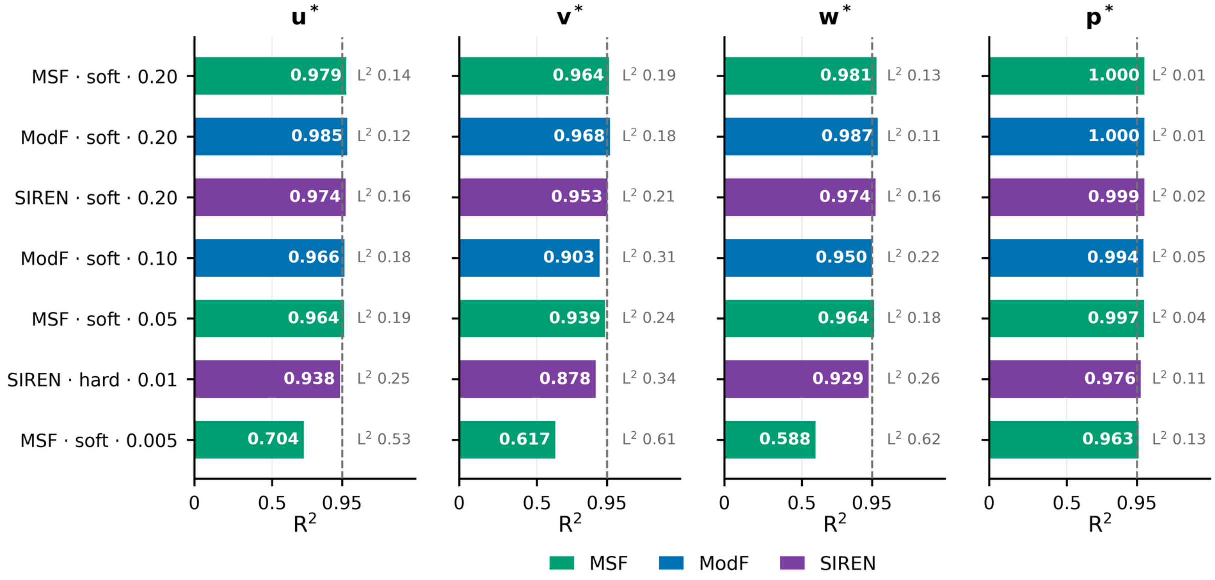


**Figure S14.** Per-field accuracy across architectures and conditions. Each panel (u, v, w, p) shows $R^2$ for seven representative configurations spanning df = 0.20 down to 0.005, colored by architecture; the corresponding relative $L^2$ is printed beside each bar. The dashed line marks $R^2$ = 0.95.

#### Supplementary Table

**Table S1.** Complete results for all 41 training runs, by architecture, inlet boundary condition, supervised-data fraction and actual training steps, with validator $R^2$ for each field at the final step. † marks the step-matched df = 0.20 runs compared in Table 3 of the main text; where two budgets exist for the same configuration both are listed. ‡ marks the data-free Multi-Scale Fourier run, trained at Re ≈ 12.5 with integral flux constraints on the combined outlets and eight internal cross-sections; every other run uses Re ≈ 251. Values were extracted directly from each run's Hydra configuration and TensorBoard event files. Main-text Table 4 is the $R^2(u)$ pivot of this table.

| Architecture | BC | df | Steps | $R^2(u)$ | $R^2(v)$ | $R^2(w)$ | $R^2(p)$ |
|---|---|---|---|---|---|---|---|
| **MLP** | soft | 0.001 | 2,200,100 | 0.699 | 0.555 | 0.653 | 0.878 |
| **MLP** | soft | 0.01 | 700,000 | 0.953 | 0.919 | 0.943 | 0.995 |
| **MLP** | soft | 0.05 | 516,100 | 0.968 | 0.945 | 0.972 | 0.999 |
| **MLP** | soft | 0.10 | 544,100 | 0.971 | 0.949 | 0.973 | 0.999 |
| **MLP** | soft | 0.20 | 500,000 | 0.972 | 0.958 | 0.980 | 1.000 |
| **MLP †** | hard | 0.20 | 300,000 | 0.968 | 0.951 | 0.976 | 0.999 |
| **Modified Fourier** | soft | 0.05 | 104,500 | 0.728 | 0.476 | 0.428 | 0.968 |
| **Modified Fourier** | soft | 0.10 | 250,000 | 0.966 | 0.903 | 0.950 | 0.994 |
| **Modified Fourier** | soft | 0.20 | 536,000 | 0.985 | 0.968 | 0.987 | 1.000 |
| **Modified Fourier** | hard | 0.01 | 23,000 | −0.021 | −0.005 | 0.031 | 0.880 |
| **Modified Fourier** | hard | 0.05 | 78,100 | 0.770 | 0.566 | 0.590 | 0.971 |
| **Modified Fourier** | hard | 0.10 | 100,000 | 0.950 | 0.857 | 0.932 | 0.990 |
| **Modified Fourier** | hard | 0.20 | 100,000 | 0.963 | 0.941 | 0.974 | 0.999 |
| **Multi-Scale Fourier** | soft | 0.005 | 2,050,900 | 0.704 | 0.617 | 0.588 | 0.963 |
| **Multi-Scale Fourier** | soft | 0.01 | 100,000 | 0.861 | 0.653 | 0.755 | 0.967 |
| **Multi-Scale Fourier** | soft | 0.05 | 100,000 | 0.964 | 0.939 | 0.964 | 0.997 |
| **Multi-Scale Fourier** | soft | 0.10 | 335,600 | 0.971 | 0.955 | 0.971 | 1.000 |
| **Multi-Scale Fourier** | soft | 0.10 | 1,000,000 | 0.965 | 0.952 | 0.960 | 1.000 |
| **Multi-Scale Fourier †** | soft | 0.20 | 300,000 | 0.979 | 0.964 | 0.981 | 1.000 |
| **Multi-Scale Fourier** | soft | 0.20 | 350,000 | 0.980 | 0.961 | 0.981 | 1.000 |
| **Multi-Scale Fourier** | hard | 0.005 | 350,000 | 0.429 | 0.360 | 0.505 | 0.947 |
| **Multi-Scale Fourier** | hard | 0.005 | 1,000,000 | 0.428 | 0.426 | 0.540 | 0.953 |
| **Multi-Scale Fourier** | hard | 0.01 | 350,000 | 0.930 | 0.868 | 0.921 | 0.986 |
| **Multi-Scale Fourier** | hard | 0.01 | 1,000,000 | 0.950 | 0.916 | 0.951 | 0.995 |
| **Multi-Scale Fourier** | hard | 0.05 | 350,000 | 0.965 | 0.943 | 0.966 | 0.999 |
| **Multi-Scale Fourier** | hard | 0.10 | 100,000 | 0.968 | 0.947 | 0.973 | 0.999 |
| **Multi-Scale Fourier** | hard | 0.10 | 350,000 | 0.971 | 0.945 | 0.966 | 1.000 |
| **Multi-Scale Fourier †** | hard | 0.20 | 300,000 | 0.978 | 0.960 | 0.981 | 1.000 |
| **Multi-Scale Fourier** | hard | 0.20 | 350,000 | 0.979 | 0.962 | 0.979 | 1.000 |

| | | | | | | | |
|---|---|---|---|---|---|---|---|
| **Multi-Scale Fourier ‡** | hard | 0 | 41,900 | −0.062 | −0.112 | −0.759 | −0.607 |
| **SIREN** | soft | 0.01 | 525,000 | 0.893 | 0.667 | 0.767 | 0.963 |
| **SIREN** | soft | 0.05 | 103,000 | 0.964 | 0.915 | 0.941 | 0.990 |
| **SIREN** | soft | 0.10 | 219,100 | 0.970 | 0.948 | 0.970 | 0.998 |
| **SIREN** | soft | 0.20 | 159,800 | 0.974 | 0.953 | 0.974 | 0.999 |
| **SIREN** | hard | 0.001 | 287,600 | −0.034 | 0.004 | 0.064 | 0.373 |
| **SIREN** | hard | 0.01 | 1,211,000 | 0.938 | 0.878 | 0.929 | 0.976 |
| **SIREN** | hard | 0.05 | 327,200 | 0.953 | 0.880 | 0.932 | 0.985 |
| **SIREN** | hard | 0.10 | 139,100 | 0.968 | 0.944 | 0.968 | 0.998 |
| **SIREN †** | hard | 0.20 | 293,100 | 0.976 | 0.957 | 0.978 | 0.999 |
| **SIREN** | hard | 0 | 48,700 | −0.019 | 0.097 | 0.207 | −0.868 |
| **SIREN** | hard | 0 | 142,700 | −0.027 | −0.029 | −0.625 | −0.180 |
